# Biosynthesis of pleuromutilin congeners using an *Aspergillus oryzae* expression platform

**DOI:** 10.1101/2022.12.03.518960

**Authors:** Fabrizio Alberti, Khairunisa Khairudin, Jonathan A. Davies, Suphattra Sangmalee, Christine L. Willis, Gary D. Foster, Andy M. Bailey

**Affiliations:** School of Biological Sciences, University of Bristol, 24 Tyndall Avenue, Bristol BS8 1TQ, UK; School of Life Sciences, University of Warwick, Gibbet Hill Road, Coventry CV4 7AL, UK; School of Chemistry, University of Bristol, Cantock’s Close, Bristol BS8 1TS, UK

## Abstract

Pleuromutilin is an antibiotic diterpenoid made by *Clitopilus passeckerianus* and related fungi, and it is the progenitor of a growing class of semi-synthetic antibiotics used in veterinary and human medicine. To harness the biotechnological potential of this natural product class, a full understanding of its biosynthetic pathway is essential. Previously, a linear pathway for pleuromutilin biosynthesis was established. Here we report two shunt pathways involving Pl-sdr and Pl-atf that were identified through the rational heterologous expression of combinations of pleuromutilin biosynthetic genes in *Aspergillus oryzae*. Three novel pleuromutilin congeners were isolated, and their antimicrobial activity was investigated, alongside that of an additional derivative produced through a semi-synthetic approach. It was observed that the absence of substituents – C-3 keto, C-11 hydroxy or C-21 keto – from the pleuromutilin core affected the antibacterial activity of pleuromutilin congeners. This study expands our knowledge on the biosynthesis of pleuromutilin and provides avenues for the development of novel pleuromutilin analogues by combining synthetic biology and synthetic chemistry.

## Introduction

Pleuromutilin is a diterpenoid antimicrobial made by *Clitopilus passeckerianus* and related basidiomycete fungi.^1,2^ Pleuromutilin’s antibacterial properties rely on the inhibition of protein synthesis by interfering with the peptidyl transferase centre (PTC) of the bacterial ribosome and subsequently preventing the formation of peptide bonds between amino acids.^3^

Numerous efforts have been made to modify the structure of pleuromutilin with the aim to improve its bioactivity and pharmacokinetic properties. Functionalisation of the C-14 side chain has led to the development of two semisynthetic derivatives used in veterinary formulations, tiamulin and valnemulin,^4^ as well as of two more derivatives, retapamulin and lefamulin, which are used in human medicine to treat skin infections and community-acquired bacterial pneumonia, respectively.^5,6^ Efforts to expand this class of antibiotics are still ongoing, including reports of new semisynthetic derivatives of pleuromutilin generated through photoinduced addition reactions at the alkene position C19-C20, which resulted in improved antimicrobial activity against Gram-positive bacteria upon introduction of an *N*-acetyl-L-cysteine side chain.^7^

Studies on the biosynthesis of pleuromutilin have been aided by the identification of the corresponding biosynthetic gene cluster in *C. passeckerianus*,^8^ which includes the seven enzyme-coding genes *Pl-ggs, Pl-cyc, Pl-p450-1, Pl-p450-2, Pl-p450-3, Pl-sdr* and *Pl-atf* (Figure 1a). Independent work from Yamane *et al*.^9^ and Alberti *et al*.^10^ led to the elucidation of the biosynthetic pathway to pleuromutilin *via* heterologous gene expression in *Aspergillus oryzae* (Figure 1b). The pathway to pleuromutilin **1** begins with Pl-ggs producing geranylgeranyl diphosphate (GGPP), while Pl-cyc is a bifunctional (di)terpene synthase^11^ that catalyses the cyclisation of GGPP to the first tricyclic intermediate 3-deoxo-11-dehydroxymutilin (also called premutilin) **2**. Two cytochrome P450s – Pl-p450-1 and Pl-p450-2 – add hydroxy groups at C-11 (producing intermediate **3**) and C-3 (producing intermediate **4**), respectively. The hydroxy group at C-3 is then oxidised to a keto by the short-chain dehydrogenase/reductase Pl-sdr, producing mutilin **5**. The acetyltransferase Pl-atf adds an acetate group to C-14, giving 14-O-acetylmutilin **6**. Lastly Pl-p450-3 oxidises C-22 on the acetate side chain to give pleuromutilin **1**. Interestingly, Yamane *et al*. ^9^ reported that Pl-p450-2 can act independently from Pl-p450-1 on the substrate **2** and leads to the accumulation of **7**, which can then be converted to **4** by Pl-p450-1 and rejoin the rest of the pathway. This suggests that both Pl-p450-1 and Pl-p450-2 have substrate flexibility. We therefore set out to investigate in the present study, if the other enzymes involved in pleuromutilin biosynthesis are also able to accept diverse substrates. Instances of alternative biosynthetic paths for secondary metabolites have been reported for natural products, such as for actinorhodin in *Streptomyces coelicolor*, which shows two alternative routes for the quinone formation by C-6 oxygenation.^12^

**Figure 1.**
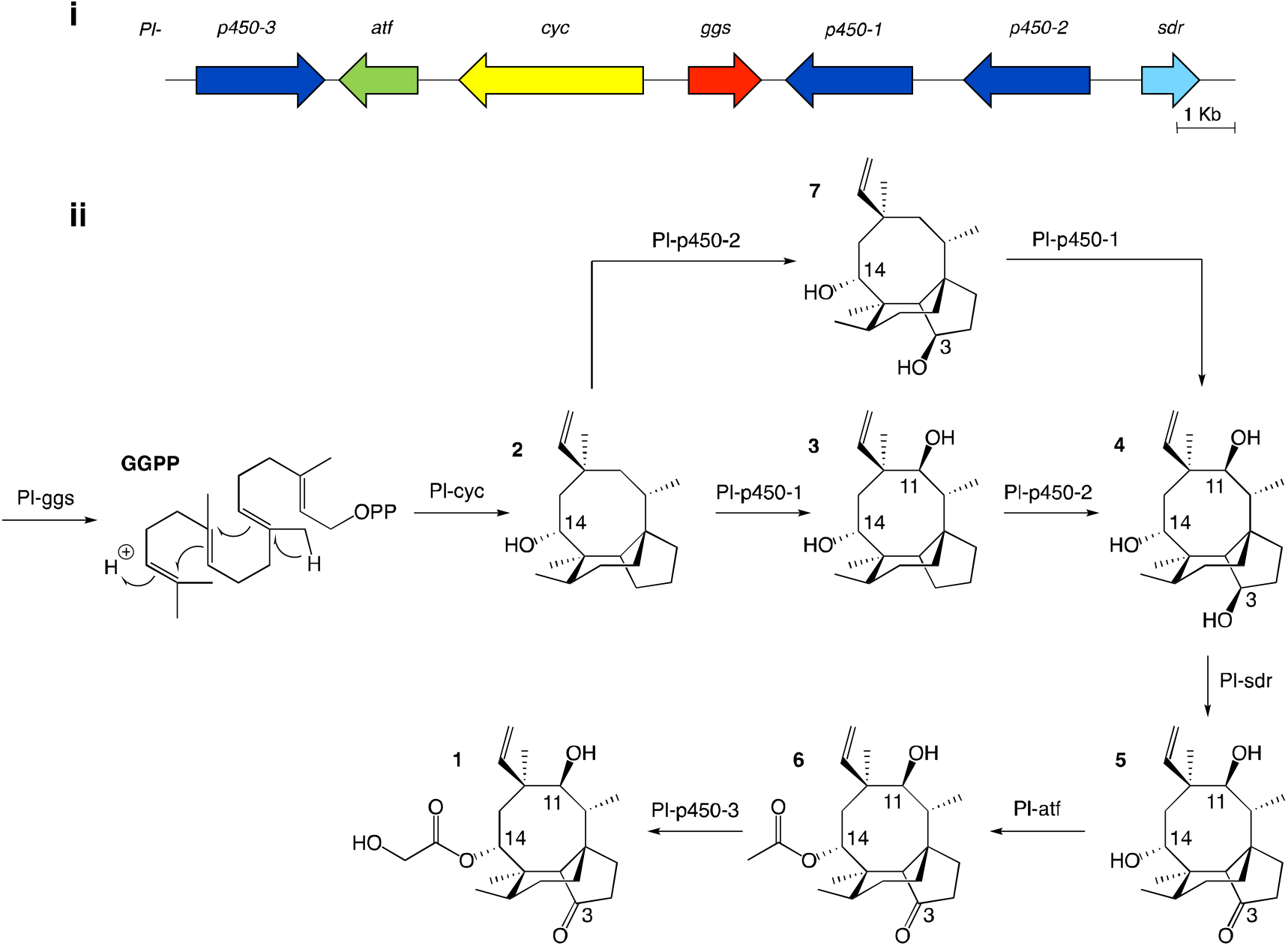
**i)** pleuromutilin biosynthetic gene cluster. **ii)** proposed biosynthetic route to pleuromutilin.^9,10^

In this study, we report the rational heterologous expression of combinations of pleuromutilin biosynthesis genes in *A. oryzae*, which led us to isolate new pleuromutilin congeners. Synthetic chemistry routes were also adopted to introduce further functionalisation, and all pleuromutilin congeners were assessed for antimicrobial activity. Results from this study aid understanding of pleuromutilin biosynthetic reactions and provide the basis for further engineering of the pleuromutilin pathway to generate new unnatural analogues.

## Results and Discussion

### Biosynthesis and isolation of novel pleuromutilin congeners

In this study, we set out to investigate the substrate tolerance of pleuromutilin biosynthetic enzymes and expand our knowledge of pleuromutilin biosynthesis. For this purpose, we used as the recipient host organism the premutilin-producing strain *A. oryzae* GC, which carries the core biosynthetic genes *Pl-ggs* and *Pl-cyc*,^10^ to further express downstream biosynthetic genes from the pleuromutilin cluster and isolate the corresponding biosynthetic products (Table 1).

**Table 1.** List of *Aspergillus oryzae* strains expressing pleuromutilin biosynthesis genes generated in this study and related compounds produced *de novo*.

| *A. oryzae* strain | Heterologous genes from *C. passeckerianus* | Compounds produced *de novo* |
| --- | --- | --- |
| GCP2 | *Pl-ggs, Pl-cyc, Pl-p450-2* | **7** |
| GCP2S | *Pl-ggs, Pl-cyc, Pl-p450-2, Pl-sdr* | **8** |
| GCP2P3SA | *Pl-ggs, Pl-cyc, Pl-p450-2, Pl-p450-3, Pl-sdr, Pl-atf* | **9** |
| GCP1P2P3A | *Pl-ggs, Pl-cyc, Pl-p450-1, Pl-p450-2, Pl-p450-3, Pl-atf* | **10** |

Firstly, we introduced *Pl-p450-2* in *A. oryzae* GC, producing strain *A. oryzae* GCP2. One major compound was produced *de novo* (see Supplementary Figure 1 for mass spectra in positive and negative ion mode) and then purified using preparative HPLC. Approximately 17 mg of white precipitate was isolated from a one-litre culture of *A. oryzae* GCP2. The compound was identified as the 3,14-diol **7** through NMR spectroscopy experiments (Figure 2, Supplementary Figures 2-6), in accordance with previous reports from Yamane *et al*.^9^

**Figure 2.**
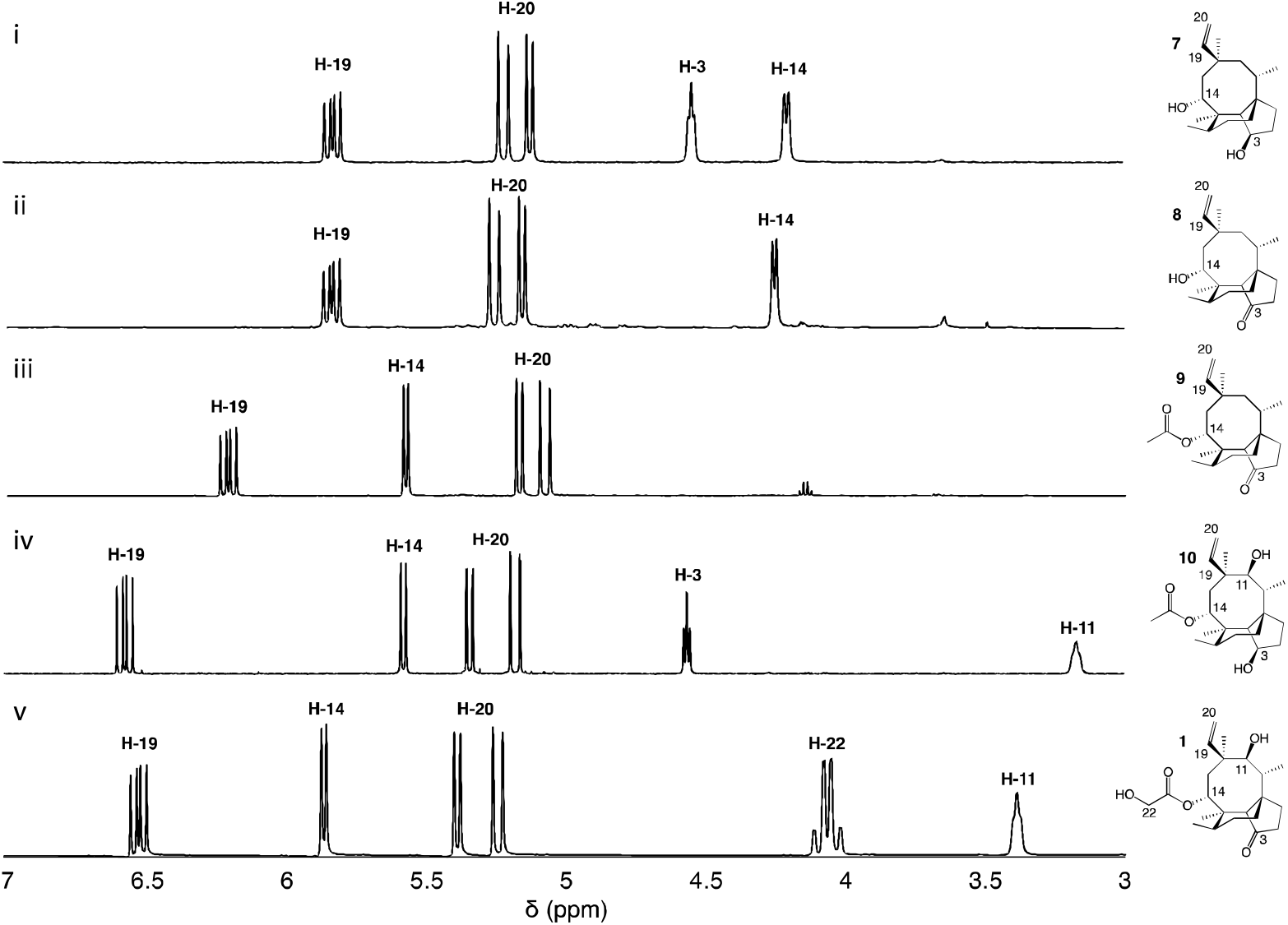
^**1**^H-NMR spectra of pleuromutilin intermediates isolated from their respective *A. oryzae* producing strains, compared to the pleuromutilin standard. The region δ 3.0–δ 7.0 ppm is shown for **i)** 3,14-diol (**7**) isolated from *A. oryzae* GCP2, **ii)** 3-keto 14-alcohol (**8**) isolated from *A. oryzae* GCP2S, **iii)** 3-keto 14-acetyl (**9**) isolated from *A. oryzae* GCP2P3SA, **iv)** 3,11-diol 14-acetyl (**10**) isolated from *A. oryzae* GCP1P2P3A, and **v)** pleuromutilin standard (**1**). Data were recorded in CDCl_3_ (500 MHz).

We next co-expressed *Pl-p450-2* and *Pl-sdr* in *A. oryzae* GC, producing strain GCP2S, and successfully purified 8.5 mg of a new product (**8**, see Supplementary Figure 7 for mass spectra in positive and negative ion mode) from a one-litre culture of the organism. The ^1^H-NMR of the product revealed similar chemical shifts as **7** except for the broad signal at δ 4.6 correlated to 3-OH, which was missing in **8**. Further extensive NMR spectroscopy studies (Figure 2, Supplementary Figures 8-12) confirmed the presence of a 3-keto as shown by the signal at δ 218.7 in the ^13^C-NMR spectrum, hence the structure of **8** was assigned as 11-dehydroxy mutilin. The isolation of **8** from *A. oryzae* GCP2S suggested that oxidation of the C-3 hydroxy group to a keto can take place independently of the hydroxylation of C-11.

The next steps in the production of mature pleuromutilin analogues involve the addition of the acetyl group on C-14 and further hydroxylation of C-22 catalysed by Pl-atf and Pl-p450-3, respectively. Thus, the expression cassettes for *Pl-p450-2* and *Pl-p450-3* alongside those for *Pl-sdr* and *Pl-atf* were transformed into *A. oryzae* GC, obtaining strain *A. oryzae* GCP2P3SA. It should be noted that this strain includes all but one – *Pl-p450-1* – pleuromutilin biosynthetic genes. HPLC analysis of crude extracts from this strain revealed a new product (**9**, see Supplementary Figure 13 for mass spectra in positive and negative ion mode). The isolation of the new product was carried out through column chromatography. The crude fungal extract from a one-litre culture of GCP2P3SA was eluted with petroleum ether/ethyl acetate (1:9) and the fractions containing the targeted compound were pooled and concentrated *in vacuo*, yielding 18 mg of white precipitate. Metabolite **9** showed similar ^1^H-NMR shift patterns as **8**, except for the signal at δ 4.25 being deshielded to δ 5.57 (Figure 2). The 2D-NMR spectroscopy experiments confirmed that this signal was correlated to H-14. Two additional carbon signals having similar chemical shifts as those observed in **6** were also observed in the ^13^C-NMR spectrum of **8**, suggesting that acetylation took place on C-14. The isolation of **9** from *A. oryzae* GCP2P3SA shows that Pl-atf can catalyse acetylation of mutilin without needing pre-hydroxylation at C-11 (Supplementary Figures 14-18). Despite the presence of *Pl-p450-3* in *A. oryzae* GCP2P3SA, no detectable C-22 hydroxylation took place. It should be noted that Pl-p450-3 shows poor catalytic activity when heterologously expressed in *A. oryzae*, which has been observed both on the native substrate 14-O-acetylmutilin,^8^ and on its congeners – as seen from the incomplete functionalisation of C22 on a C1-C2 unsaturated 14-O-acetylmutilin analogue.^10^ Nevertheless, based on these results, we propose that intermediate **7** can be derivatised at positions C-3 and C-14 leading to two previously unreported pleuromutilin congeners, **8** and **9**, through the catalytic activity of Pl-sdr and Pl-atf, respectively.

We next tested the impact of omitting *Pl-sdr* from the gene cluster. We therefore generated strain *A. oryzae* GCP1P2P3A, which included the three pathway cytochrome P450s and *Pl-atf* in the *A. oryzae* GC background. The *A. oryzae* GCP1P2P3A strain accumulated a novel compound (**10**, see Supplementary Figure 19 for mass spectra in positive and negative ion mode), which was purified through preparative HPLC yielding 20 mg of white precipitate. Analysis of the NMR spectroscopy data for **10** revealed the presence of two hydroxy groups. From the ^13^C-NMR spectrum (Supplementary Figures 20-24), it appeared that this molecule would be related to the previously characterized **4**, though bearing two additional carbons. The chemical shifts of these two carbons matched with the shift pattern of C-21 and C-22 from **6** and the ^1^H-NMR data also demonstrated that the C-14 had incorporated an acetyl group, giving **10**. The isolation of **10** from *A. oryzae* GCP1P2P3A proved that the C-14 acetylation reaction catalysed by Pl-atf can occur not only when there is a keto group at C-3 – as it is the case for mutilin **5** in the main pathway – but also in the presence of a hydroxy group at the same position – as it is the case for **4**. As with metabolite **9**, the acetate group on C-14 of **10** was not oxidized despite *Pl-p450-3* being expressed in *A. oryzae* GCP1P2P3A. Isolation of this shunt product suggested that the acetylation catalysed by Pl-atf could take place independently from the oxidation of the C-3 hydroxy to keto. Overall, it appeared that when the activity of Pl-p450-1 and Pl-sdr were omitted, alternative pathways could be followed bypassing the catalytic reactions of the missing enzymes and leading to shunt products. These results suggest that some of the pleuromutilin biosynthetic enzymes possess relaxed substrate tolerance, which makes them useful as potential biocatalysts to generate novel pleuromutilin analogues. Using a similar approach to the one we adopted here, analogues of the final biosynthetic products from other pathways have been obtained in other studies. For instance, the biosynthetic enzymes involved in the production of the communesins show a good degree of substrate tolerance to make analogues of the final pathway products, as Lin *et al*.^13^ were able to generate communesin C, E, F and J, analogous to communesin A and B, from blocked mutants lacking the methyltransferase and epoxidase enzymes CnsE and CnsJ.

### Production of metabolites 9 and 10 through feeding experiments

The rational heterologous expression of all genes of the pleuromutilin cluster minus *Pl-p450-1* or *Pl-sdr* showed production of new metabolites, **9** and **10**. However, the final hydroxylation of C-22 catalysed by Pl-p450-3 did not take place on either of these two shunt products. To investigate this reaction further, we performed feeding studies with 11-dehydroxy mutilin **8** and triol **4** on *A. oryzae* AP3, which harbours *Pl-p450-3* and *Pl-atf* and was previously shown to promote conversion of **5** into **1**.^10^ Metabolites **8** and **4** were isolated from strains *A. oryzae* GCP2S and GCP1P2, respectively. *A. oryzae* AP3 was cultured in the presence of 100 mg l^-1^ of **8** and **4** in separate experiments. Metabolite analyses of the fed cultures confirmed the conversion of **8** to **9** (Supplementary Figure 25) and of **4** to **10** (Supplementary Figure 26), however no congener with hydroxylated C-22 could be detected in either assay, in accordance with our previous observations of metabolites production in strains GCP2P3SA and GCP1P2P3A.

### Antimicrobial activity testing of novel pleuromutilin congeners

The new shunt products isolated in this work were assayed for antimicrobial activity on *Bacillus subtilis* ATCC 6633 and compared with the intermediates known to be part of the main pathway, following the same procedure reported by Alberti *et al*.^10^ Pleuromutilin **1** showed the highest antimicrobial activity, followed by 14-O-acetylmutilin **6** (Figure 3). The small difference in bioactivity between **1** and **6** is in accordance with observations that the C-14 extension of pleuromutilin antibiotics is responsible for only minor hydrophobic interactions with bacterial ribosomal nucleotides.^3^ Conversely, the loss of the keto group at C-21 had a considerable impact on bioactivity, as can be seen in the reduced inhibitory activity of **3, 4, 7** and **10**. This is in line with observations of the interaction of pleuromutilins with the ribosomal nucleotides, in which the C-21 keto group is known to form a network of hydrogen bonds with G2061 of the 23S RNA domain V.^3^ Mutilin **5** shows approximately half the activity of pleuromutilin while earlier intermediates with reduction of keto to hydroxy group or absence of the hydroxy group from C-3, **4** and **3** respectively, resulted in almost complete loss of activity. This is an agreement with previous studies showing that mutilin and other derivatives with a free C-14 hydroxy group were inactive.^14^ Absence of the C-11 hydroxy group – known to form hydrogen bonding with the G2505 phosphate of the 23S RNA domain V^3^ – also resulted in a considerable decline in bioactivity for **7** and **8**, although **8** showed a slightly stronger inhibition putatively due to the presence of a keto group at C-3. Similar to pleuromutilin intermediates, a decline in antimicrobial activity was also observed in the shunt products **10** and **9** when these are compared to 14-O-acetylmutilin **6**, highlighting the importance of the C-3 keto and the C-11 hydroxy groups, respectively, for bioactivity.

**Figure 3.**
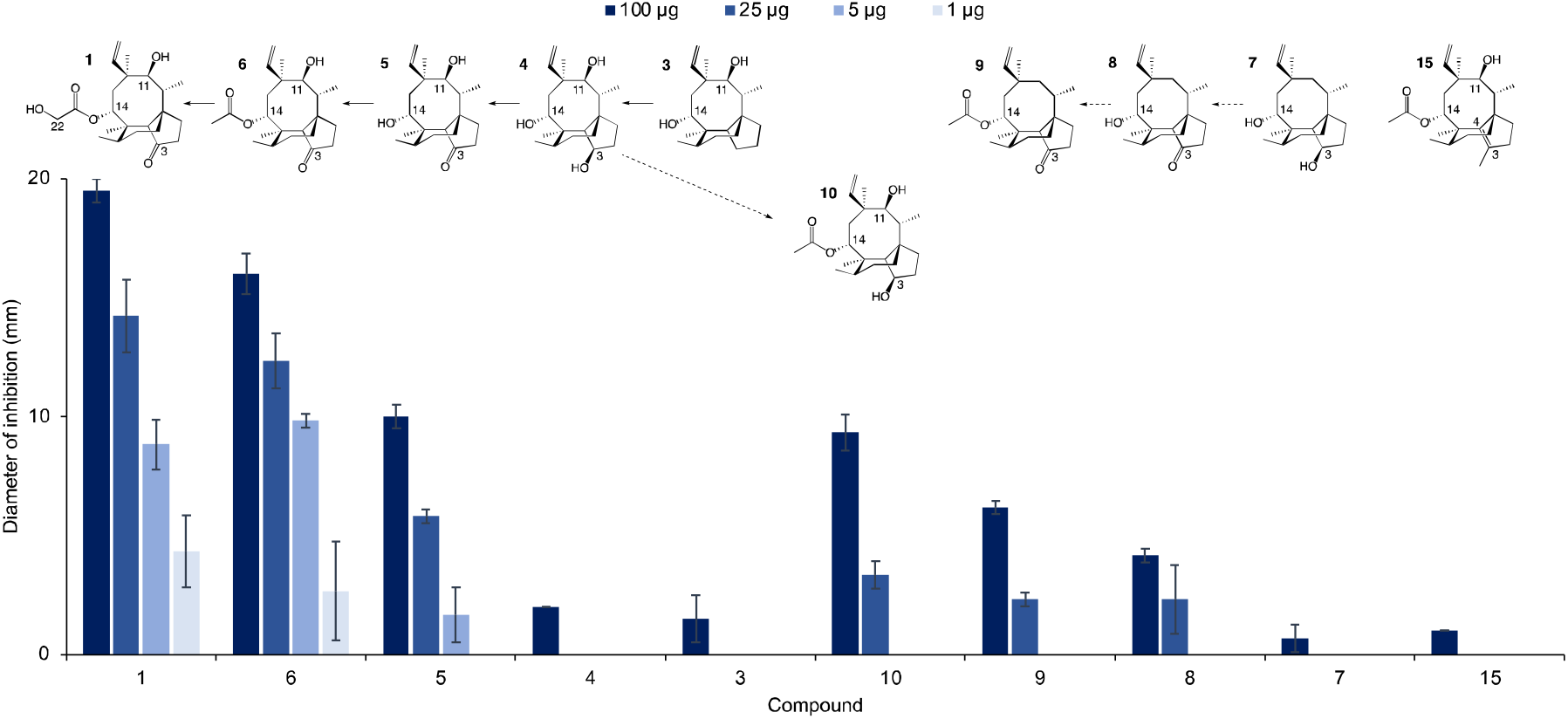
Antimicrobial activity of pleuromutilin (**1**), its intermediates (**3-6**), shunts (**7-10**) and semisynthetic derivative (**15**), against *B. subtilis*. Reduction of the antimicrobial activity is observed upon loss of specific substituent groups at C-14, C-3, and C-11 of the pleuromutilin core. The chemical structure of each metabolite is shown above the graph. Solid arrows indicate that biosynthetic reactions are part of the main pathway, whereas dashed arrows denote reactions involved in shunt pathways.

**Figure 4.**
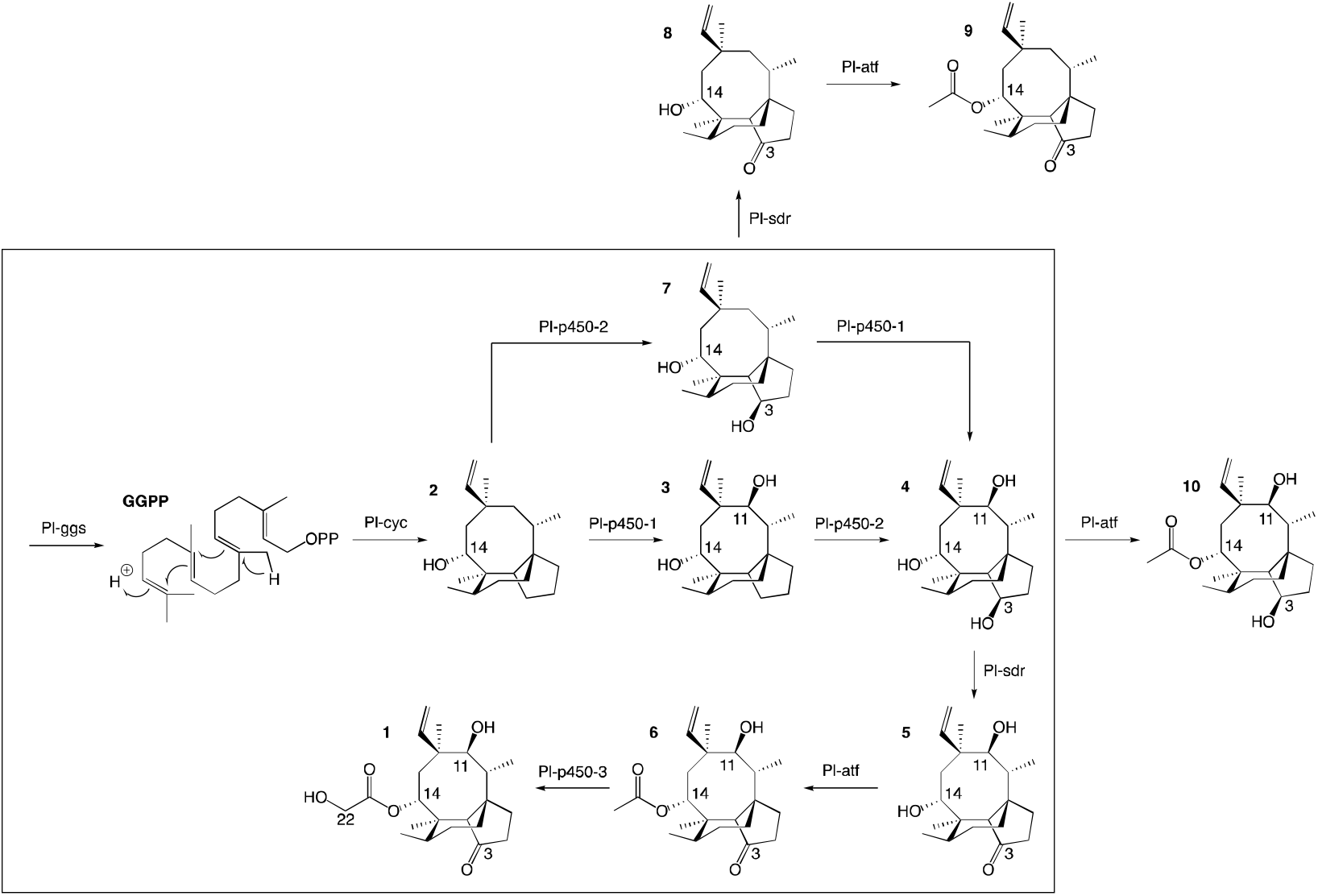
Revised pleuromutilin biosynthetic pathway. The main pathway is in the box, the shunt products are outside.

Given the importance of the C-3 keto group in bioactivity, we decided to investigate whether substitution of this group with other functional groups would lead to an altered antimicrobial activity. Furthermore, the five-membered ring of pleuromutilin antibiotics is prone to attack by liver cytochrome P450s, which leads to their rapid metabolism.^15^ Therefore, their functionalisation may result in pleuromutilin derivatives with longer half-life or even altered bioactivity, as shown recently by the development of an anti-idiopathic pulmonary fibrosis lead compound through the functionalisation of the five-membered ring of pleuromutilin.^16^ In this study, we endeavoured to functionalise the five-membered ring by replacing the C-3 keto of pleuromutilin with a methyl group coupled with C3-C4 unsaturation. Hydrolysis of commercially available tiamulin hydrogen fumarate was performed to produce mutilin **5** (see Supplementary Figures 27-28 for ^1^H-NMR and ^13^C-NMR spectra, and Supplementary Methods for reaction conditions). Mutilin was used to prepare TMS-mutilin **12** (see Supplementary Figures 29-30 for ^1^H-NMR and ^13^C-NMR spectra, and Supplementary Methods for reaction conditions), which was then used to make a C-3 methyl, C3-C4 unsaturated analogue **14** (see Supplementary Figures 31-32 for ^1^H-NMR and ^13^C-NMR spectra, and Supplementary Methods for reaction conditions). Analogue **14** was fed to *A. oryzae* AP3 at a concentration of 100 mg l^-1^ for further functionalisation of the C-14 side chain. Successful acetylation by Pl-atf was observed, leading to the new semi-synthetic analogue **15**, which was purified using a combination of column chromatography (10-100% ethyl acetate in petrol) and preparative HPLC, yielding 23 mg of pure compound (see Supplementary Figures 33-34 for ^1^H-NMR and ^13^C-NMR spectra). Once again, no congener with hydroxylated C-22 could be detected, as previously observed when attempting to produce pleuromutilin analogues starting from **4** and **8**. The semi-synthetic compound **15** was assayed for antimicrobial activity against *B. subtilis*. Compared to the closest naturally occurring intermediate **6**, a considerable decrease in activity was observed when the five-membered ring was functionalised with a C-3 methyl and a C3-C4 double bond (Figure 3). Thus, from our study, it can be concluded that retaining the three naturally occurring pleuromutilin functional groups – C-3 keto, C-11 hydroxy and C-21 keto – is important to maintain the bioactivity of the pleuromutilin analogues.

## Conclusions

A more detailed picture of the biosynthetic pathway for pleuromutilin was obtained in this work, showing that biosynthetic genes can be expressed in different combinations to produce pleuromutilin congeners that have not been reported in *C. passeckerianus*. By omitting selected biosynthetic genes, we were able to isolate three novel pleuromutilin congeners, **8, 9** and **10**. Overall, it appeared that when the activity of Pl-p450-1 and Pl-sdr were omitted, alternative pathways could be followed bypassing the catalytic reactions of the missing enzymes and leading to shunt products. These results suggest that some of the pleuromutilin biosynthetic enzymes possess relaxed substrate tolerance, which makes them useful as potential biocatalysts to generate novel pleuromutilin analogues. Heterologous gene expression in *A. oryzae* proved to be a flexible approach for developing modified pleuromutilin analogues. This strategy, when combined with synthetic endeavours, opens to the possibility of generating new semi-synthetic pleuromutilins.

## Materials and methods

### Reagents, strains and conditions for growth of microorganisms

Chemicals and media ingredients used in this study were obtained from Sigma, Fisher, Oxoid, ForMedium, Melford, Bioline, Thermo Scientific or VWR unless otherwise stated. Media were prepared and sterilized by autoclaving using a standard programme at 121 °C for 15 minutes. Deionized water was used for all solutions unless stated.

### Growth condition for fungal and bacterial strains

The host strain used for heterologous expression, *A. oryzae* NSAR1 (genotype niaD-, sC-, ΔargB, adeA-)^17^ was maintained on MEA plates with appropriate supplements (15 g l^-1^ malt extract, 1.5 g l^-1^ arginine, 1.5 g l^-1^ methionine, 0.1 g l^-1^ adenine, 2 g l^-1^ ammonium sulphate, 15 g l^-1^ agar) at 28°C. The yeast strain used for homologous recombination-based construction of plasmids, *S. cerevisiae* BY4742 (genotype MATα, his3Δ1, leu2Δ0, lys2Δ0, ura3Δ0), was maintained on YPAD plates (10 g l^-1^ yeast extract, 20 g l^-1^ bactopeptone, 20 g l^-1^ D-glucose, 40 mg l^-1^ adenine hemisulfate 15 g l^-1^ agar). Propagation of plasmids was performed in *E. coli* One Shot® ccdB SurvivalTM 2 T1R competent cells (Life Technologies), which was grown in LB agar plates (10 g l^-1^ NaCl, 10 g l^-1^ tryptone, 5 g l^-1^ yeast extract, pH 7) at 37°C.

### Construction of expression vectors

To construct fungal expression vector containing intron-free genes of the pleuromutilin cluster, three pTYGS^18^ plasmids with either arginine, adenine or basta selectable marker were used as backbones (plasmid maps are shown in Supplementary Methods 1-4). The assembly of expression vectors pTYGSargGC, pTYGSadeP1P2P3, pTYGSadeP3, pTYGSbarAS, pTYGSbarA and pTYGSbarS was reported in Alberti *et al*.^10^ The construction of expression vectors pTYGSadeP2P3 and pTYGSadeP2 was carried out through yeast-based homologous recombinant as described by Ma *et al*.^19^ using PCR-amplified genes of interest with Phusion high-fidelity DNA polymerase according to the manufacturer’s instructions (details on all plasmid components are reported in Supplementary Table 1, plasmid maps are represented in Supplementary Figure 35). Primers used for amplification of the genes of interest are listed in Supplementary Table 2.

### Transformation of *A. oryzae* NSAR1

Protoplast−polyethylene glycol method adapted from Halo *et al*.^20^ was used to transform *A. oryzae* NSAR1 (see Supplementary Table 3 for a list of strains generated in this study). The strain was firstly transformed with vector pTYGSargGC (which included *Pl-ggs* and *Pl-cyc*) to generate strain GC (producer of **2**). Further transformation of GC with pTYGSadeP2 (containing *Pl-p450-2*) achieved production of strain GCP2 (producer of **7**). Likewise, co-transformation of GC with pTYGSadeP2 (containing *Pl-p450-2*) and pTYGSbarS produced strain GCP2S (producer of **8**). To construct GCP2P3SA (producer of **9**), a combination of pTYGSadeP2P3 (containing *Pl-p450-2* and *Pl-p450-3*) and pTYGSbarSA (containing *Pl-sdr* and *Pl-atf*) was introduced into GC, whereas a combination of pTYGSadeP1P2P3 (containing the three *Pl-p450s*) and pTYGSbarA (containing *Pl-atf*) was used to transform strain GC to obtain strain GCP1P2P3A (producer of **10**). Strain AP3, used for feeding experiments with **4** and **8**, was generated by transforming *A. oryzae* NSAR1 with a combination of pTYGSadeP3 (containing *Pl-p450-3*) and pTYGSbarA (containing *Pl-atf*), as previously reported.^10^ *A. oryzae* transformant strains were screened by PCR for integration of the genes of interest using DreamTaq Green PCR Master Mix (Thermo Scientific) in combination with the screening primers reported in Supplementary Table 2.

### Metabolite analysis of *A. oryzae* transformants

*A. oryzae* transformant strains were analysed for production of metabolites through HPLC-MS. Each strain was grown in 100 ml of CMP medium (35 g l^−1^ Czapek-dox liquid, 20 g l^−1^ maltose, 10 g l^−1^ peptone) at 28 °C for 10 days prior to performing metabolite extractions with ethyl acetate, followed by concentration of the crude extract *in vacuo* methanol. Crude extracts were dissolved in methanol and analysed by HPLC-MS including a Waters 2767 HPLC system with a Waters 2545 pump, a Phenomenex LUNA column (2.6 μ, C18, 100 AC, 4.6 × 100 mm) and a Phenomenex Security Guard precolumn (Luna C5 300 AC) for reverse-phase chromatography. UV absorbance was detected between 200 – 400 nm through a Waters 2998 diode array detector while a mass range between 150 and 800 Da in positive and negative ion mode was scanned by the Waters Quattro Micro spectrometer. Chromatography was achieved using a gradient of solvents (A, HPLC grade H_2_O containing 0.05% formic acid; B, HPLC grade CH_3_CN containing 0.045% formic acid) with the following programme: 0 minutes, 20% B; 15 minutes, 90% B; 16 minutes 95% B; 17 minutes 95% B; 18 minutes 10% B, 20 minutes 10% B. The flow rate was set at 1 ml min^-1^.

### Isolation and purification of metabolites of interest

Large scale metabolite extractions from 1-litre cultures of chosen fungal strains in CMP medium were performed with the same conditions as described above. Purification of compound **9** was carried out through column chromatography by passing the concentrated crude extract through a packed silica gel 60 (Merck) in a column. An appropriate gradient solvent system consisting of a mixture of ethyl acetate:petroleum ether (9:1 to 7:3, v/v) was used to elute the metabolites into fractions. Fractions with the targeted compound were pooled in a round bottom flask and dried *in vacuo*. Isolation of other shunt products was performed through preparative HPLC with a reverse-phase Phenomenex LUNA column (5μ, C18, 100 A□, 10 × 250 mm) using a flow rate at 16 ml min^-1^. The mobile phase was a gradient mixture of acetonitrile and formic acid in water following the elution programme: 0 minutes, 5% B; 1 minutes, 5% B; 2 minutes, 40% B; 15 minutes 90% B; 17 minutes 95% B; 18 minutes 5% B, 20 minutes 5% B (A, HPLC grade H_2_O containing 0.05% formic acid; B, HPLC grade CH_3_CN containing 0.045% formic acid). The eluent was monitored by UV detection at 200-400 nm and mass range 100-600 Da. Targeted compounds were isolated and collected in glass vials prior to being pooled and concentrated *in vacuo*.

### NMR spectroscopy analysis of purified metabolites

Nuclear Magnetic Resonance (NMR) spectroscopy was used routinely to characterise purified metabolites. Dried compounds were dissolved in CDCl_3_ and characterised by NMR spectroscopy, which was conducted either on an Agilent VNMRS500 spectrometer (^1^H NMR at 500 MHz and ^13^C NMR at 125 MHz) or on an Agilent V400-MR spectrometer (^1^H NMR at 400 MHz and ^13^C NMR at 100 MHz). Chemical shifts were recorded in parts per million (ppm) and coupling constant (J) in Hz.

### Assessing antimicrobial activity of purified metabolites

The purified compounds were tested against *Bacillus subtilis* to study antimicrobial activity. A 1×10^9^ ml^-1^ spore suspension of *B. subtilis* with 4% 2,3,5-Triphenyl-2H-tetrazolium chloride (TTC) was overlaid onto TSA agar (30 g l^-1^ tryptic soy, 15 g l^-1^ agar). Purified compounds in different amounts of 100, 25, 5 and 1 μg were each released onto individual sterile 6-mm paper discs and placed on the bacterial lawn. All plates were incubated overnight at 28 °C, following which the growth inhibition zones were measured.

## Author contributions

GDF and AMB coordinated the project. FA, KK and SS performed the heterologous expression and characterisation of metabolites produced *de novo*, under the supervision of AMB and GDF. JAD performed chemical synthesis under the supervision of CLW. KK and FA drafted the manuscript with edits and contributions from AMB and GDF.

## Acknowledgments

Dr Colin Lazarus is thanked for providing expression vectors for the transformation of *A. oryzae*; the NMR Spectroscopy and MS facilities and teams of the University of Bristol are thanked for data collection and helpful discussion. KK was supported by a scholarship from Majlis Amanah Rakyat. JAD was supported by a scholarship from the EPSRC, Bristol Chemical Sciences Centre for Doctoral Training (EP/L015366/1). FA would like to acknowledge current funding from UKRI through a Future Leaders Fellowship (MR/V022334/1).

