## Supplementary Information for "Biosynthesis of pleuromutilin congeners using an *Aspergillus oryzae* expression platform"

### 1. Supplementary Data

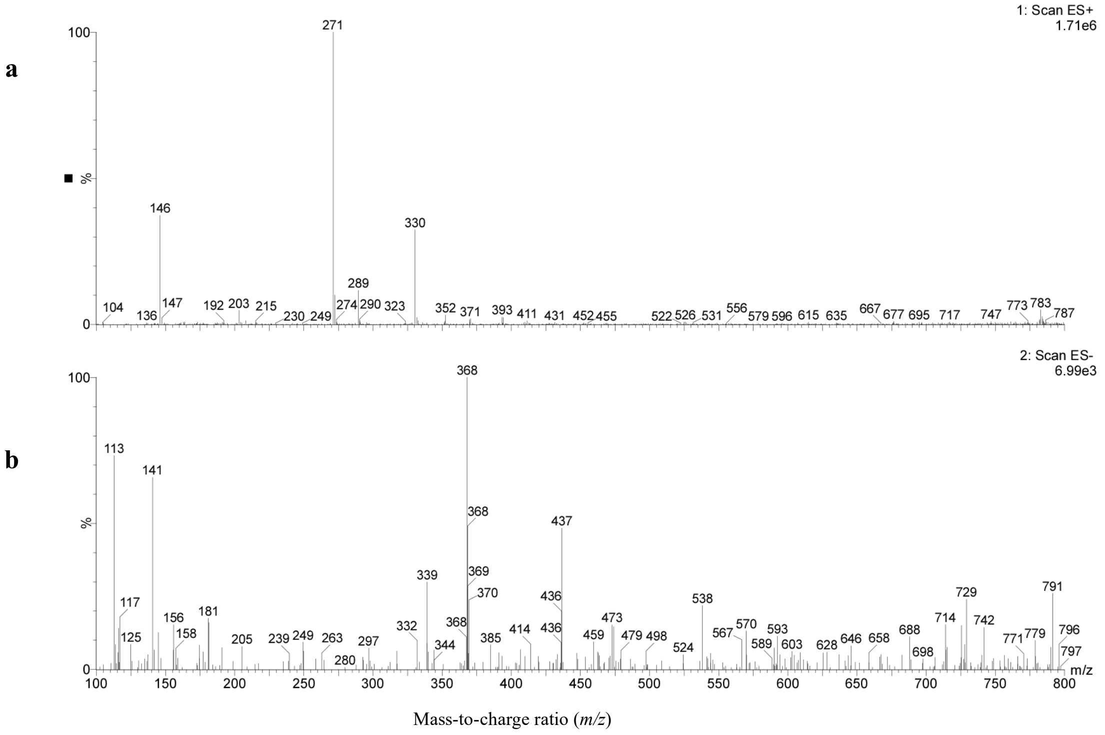

**Supplementary Figure 1.** Mass spectra in positive (a) and negative (b) ion mode of **7.**

**NMR data assignment for 7 in $CDCl_3$**

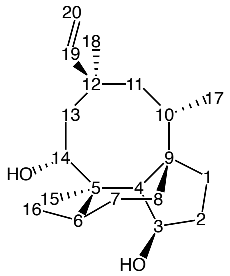

$\delta_H$ (500 MHz, $CDCl_3$) 5.82 (1H, dd, *J* 17.8, 11.1, 19-H), 5.21 (1H, dd, *J* 17.8, 1.3, 20-*H*H), 5.12 (1H, dd, *J* 11.1, 1.3, 20-H*H*), 4.54 (1H, t, *J* 5.8, 3-H), 4.20 (1H, d, *J* 8.0, 14-H), 2.25 (1H, ddp, *J* 11.4, 7.1, 3.6, 6-H), 1.97 (1H, m, 8-*H*H), 1.90 (1H, m, 13-*H*H), 1.80 (1H, p, *J* 7.0, 10-H), 1.72 (1H, d, *J* 4.4, 2-*H*H), 1.58 (1H, m, 4-H), 1.62-1.57 (2H, m, 2-H*H*, 8-H*H*), 1.54-1.47 (3H, m, 1-*H*H, 7-*H*H, 13-H*H*), 1.40-1.33 (3H, m, 1-H*H*, 7-H*H*, 11-*H*H), 1.16 (1H, s, 11-H*H*), 1.14 (3H, s, 15-$H_3$), 0.96 (3H, d, *J* 7.4, 16-$H_3$), 0.94 (3H, s, 18-$H_3$), 0.84 (3H, d, *J* 6.8, 17-$H_3$). $\delta_C$ (125 MHz, $CDCl_3$) 147.1 (C-19), 113.1 (C-20), 78.1 (C-3), 69.0 (C-14), 52.0 (C-4), 46.5 (C-13), 46.1 (C-9), 44.2 (C-11), 42.1 (C-5), 40.6 (C-12), 37.0 (C-6), 34.2 (C-8), 33.1 (C-18), 32.6 (C-2), 32.0 (C-1), 31.3 (C-10), 28.3 (C-7), 20.9 (C-17), 18.9 (C-16), 16.5 (C-15). HRMS (ESI) calc. for $C_{20}H_{34}O_2Na^+$ 329.2451. Found 329.2455.

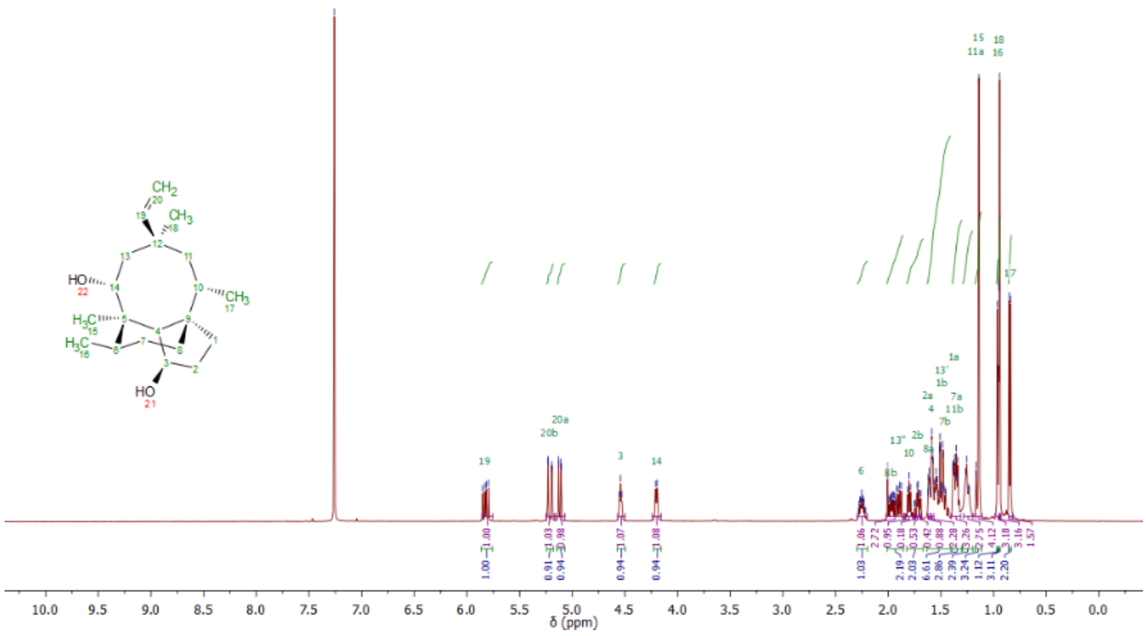

**Supplementary Figure 2.** $^{1}$H-NMR spectrum of **7** in $CDCl_3$ (500 MHz).

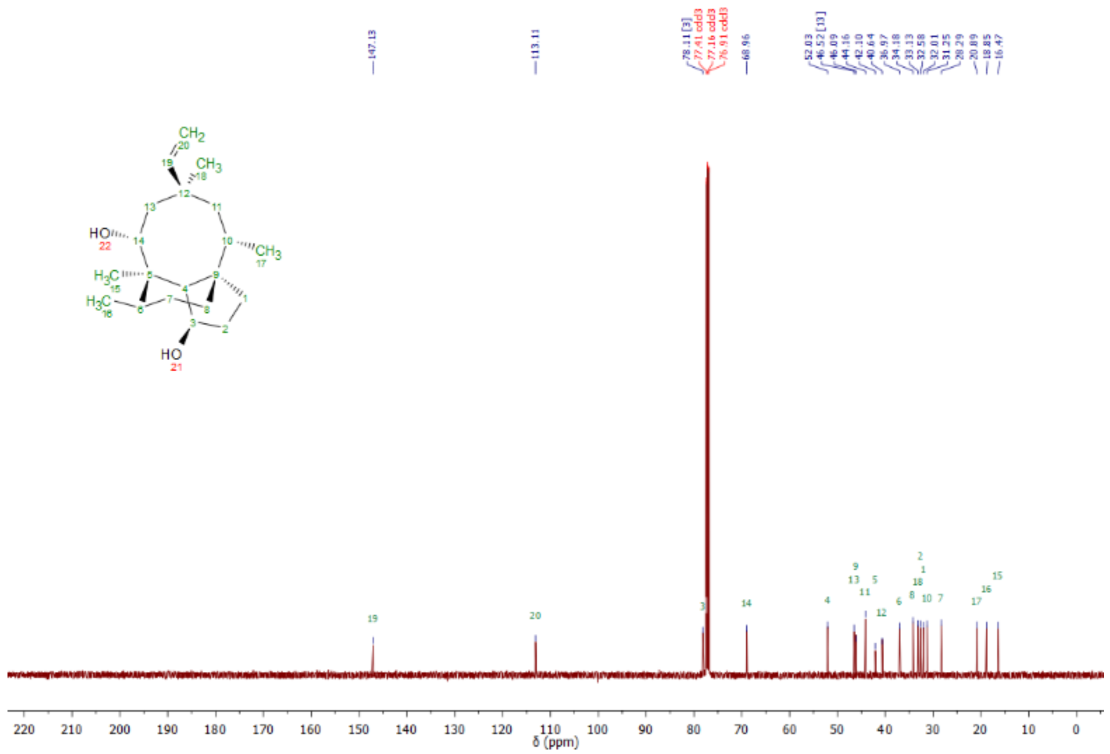

**Supplementary Figure 3.** $^{13}$C-NMR spectrum of **7** in $CDCl_3$ (125 MHz).

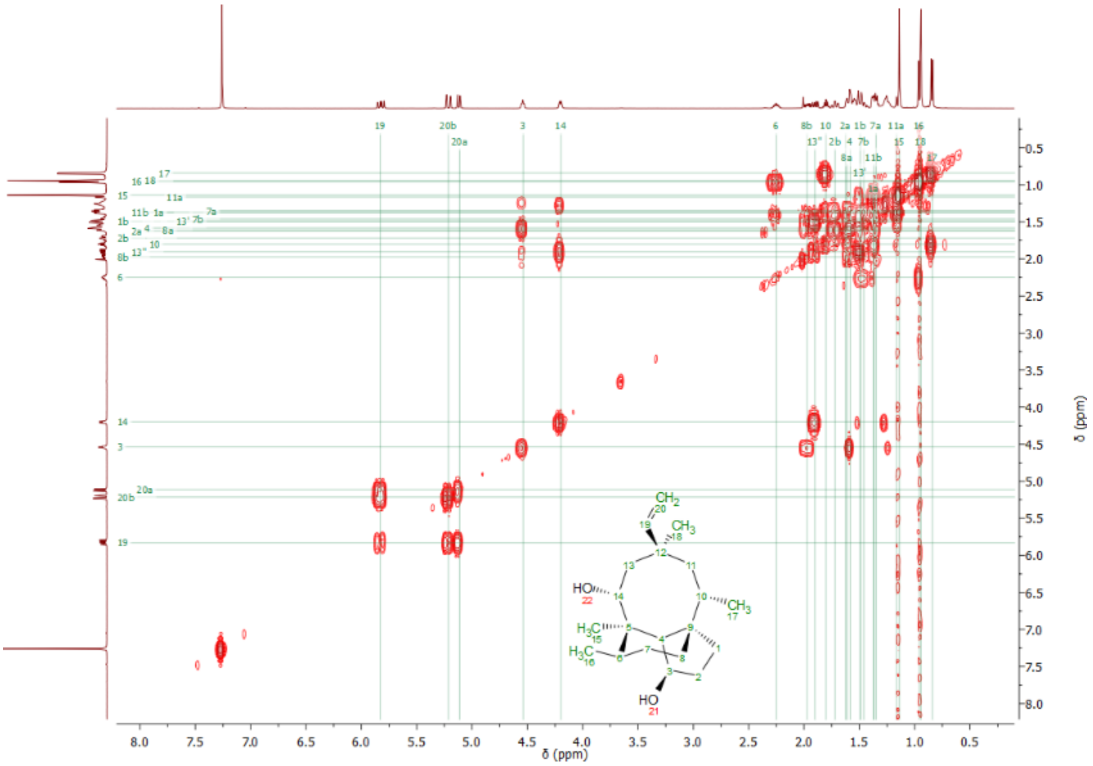

**Supplementary Figure 4.** COSY spectrum of **7** in $CDCl_3$ (500 MHz).

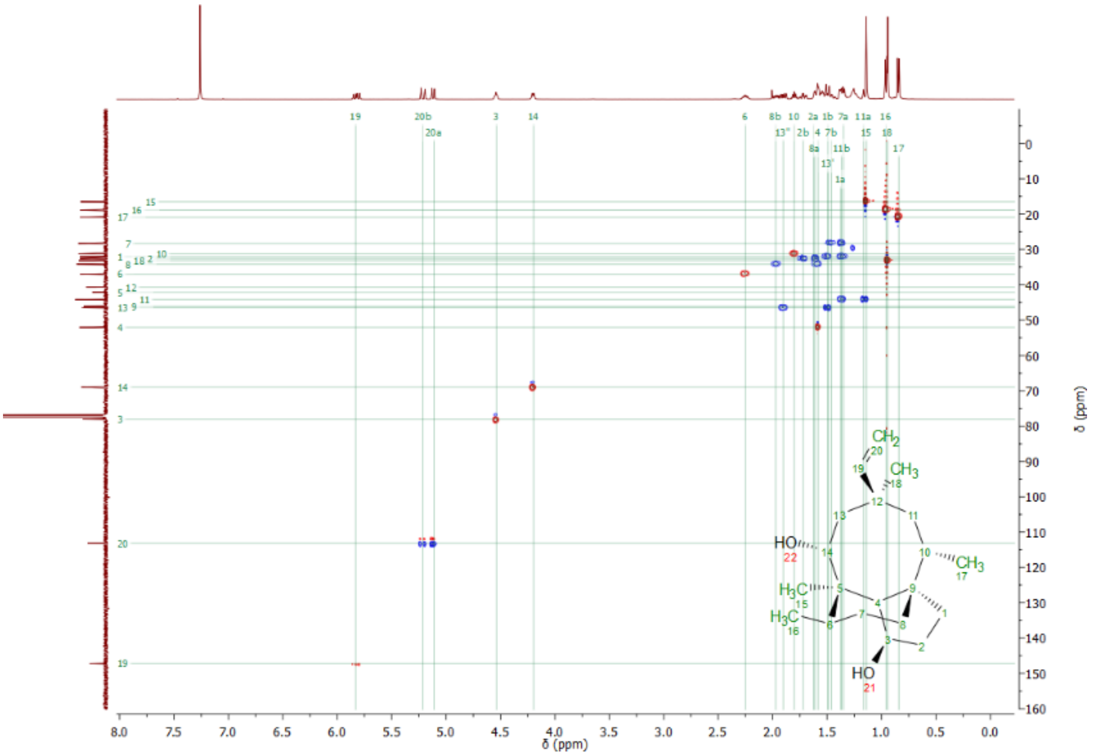

**Supplementary Figure 5.** HSQC spectrum of **7** in $CDCl_3$ (500 MHz).

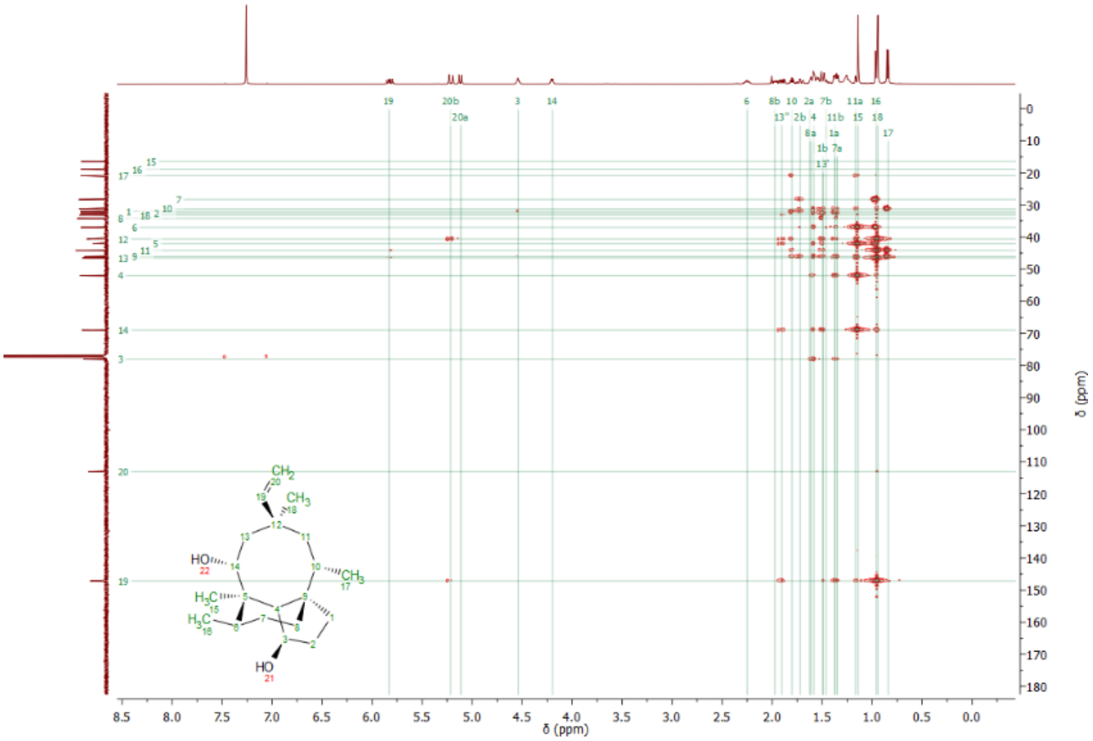

**Supplementary Figure 6.** HMBC spectrum of **7** in $CDCl_3$ (500 MHz).

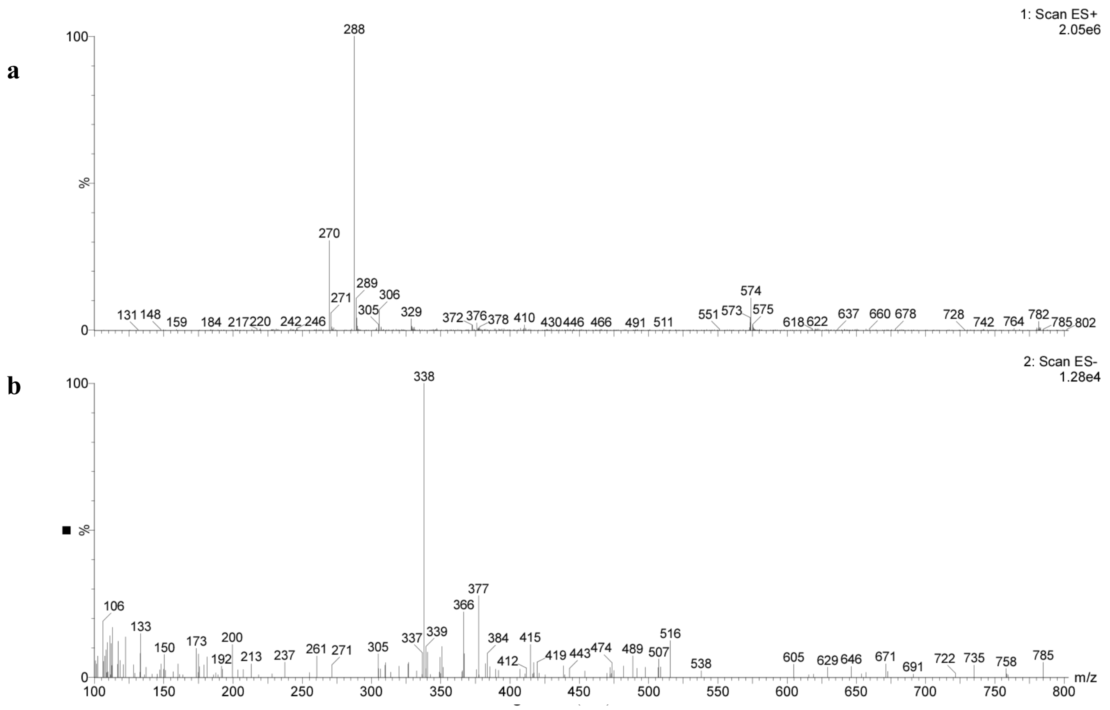

**Supplementary Figure 7.** Mass spectra in positive (a) and negative (b) ion mode of **8**.

**NMR data assignment of 8 in $CDCl_3$**

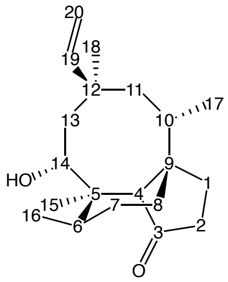

$\delta_H$ (500 MHz, $CDCl_3$) 5.84 (1H, dd, *J* 17.8, 11.0, 19-H), 5.25 (1H, d, *J* 17.8, 20-*H*H), 5.15 (1H, d, *J* 11.0, 20-H*H*), 4.25 (1H, d, *J* 7.2, 14-H), 2.20 (2H, m, 2-$H_2$), 2.10 (1H, s, 4-H), 1.97 (1H, m, 10-H), 1.86 (1H, dd, *J* 15.4, 7.3, 13-*H*H), 1.67 (1H, m, 6-H), 1.64 (1H, m, 8-*H*H), 1.61-1.47 (4H, m, 1-*H*H, 7-*H*H, 11-*H*H, 13-H*H*), 1.40 (1H, m, 1-H*H*), 1.38 (1H, m, 7-H*H*), 1.37 (3H, s, 15-$H_3$), 1.29 (1H, m, 11-H*H*), 1.12 (1H, m, 8-H*H*), 0.96 (6H, m, 16-$H_3$, 18-$H_3$), 0.90 (3H, d, *J* 6.9, 17-$H_3$). $\delta_C$ (125 MHz, $CDCl_3$) 218.7 (C-3),146.1 (C-19), 113.4 (C-20), 67.6 (C-14), 59.5 (C-4), 46.5 (C-13), 45.5 (C-12), 43.4 (C-11), 42.2 (C-5), 39.9 (C-9), 37.1 (C-6), 34.5 (C-2), 32.7 (C-18), 31.5 (C-10), 30.1 (C-8), 27.2 (C-7), 25.1 (C-1), 19.7 (C-17), 18.2 (C-16), 13.5 (C-15). HRMS (ESI) calc. for $C_{20}H_{32}O_2Na^+$ 327.2294510. Found 327.228310.

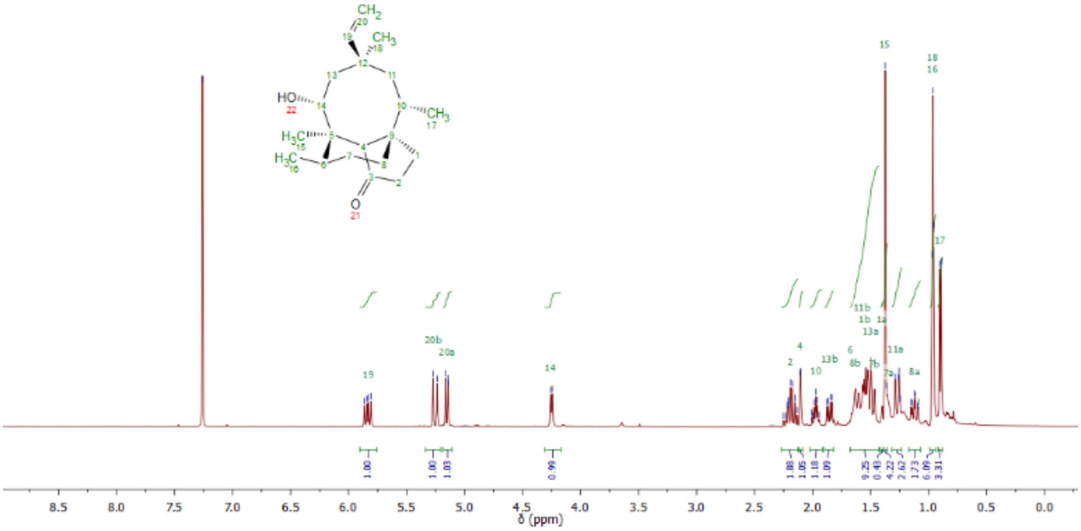

**Supplementary Figure 8.** $^{1}$H-NMR spectrum of **8** in $CDCl_3$ (500 MHz).

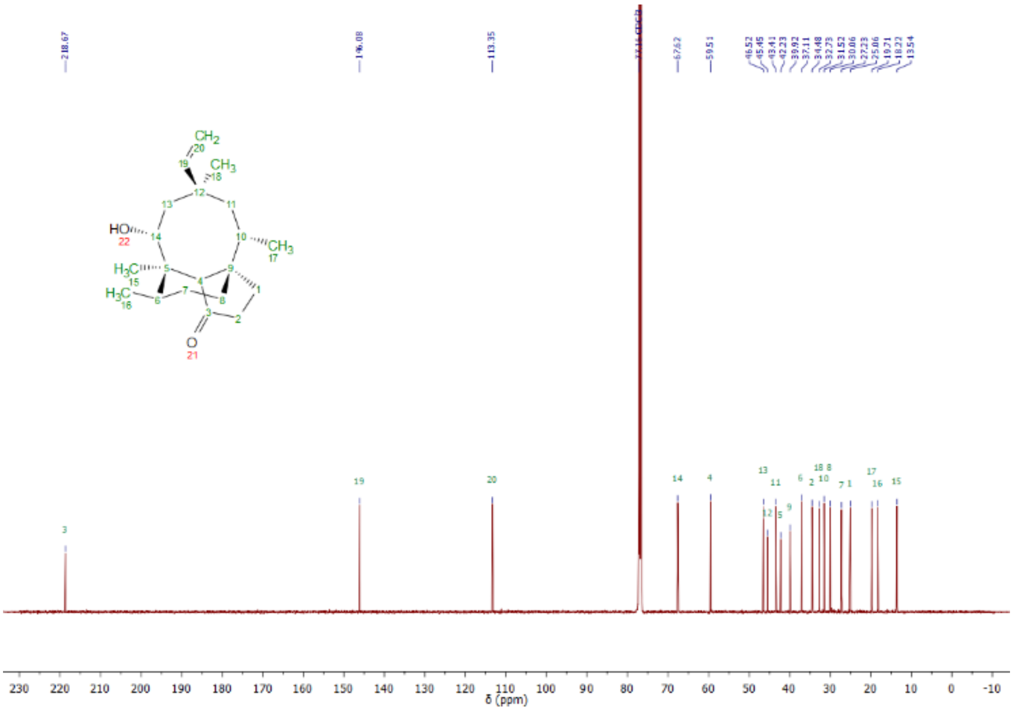

**Supplementary Figure 9.** $^{13}$C-NMR spectrum of **8** in $CDCl_3$ (125 MHz).

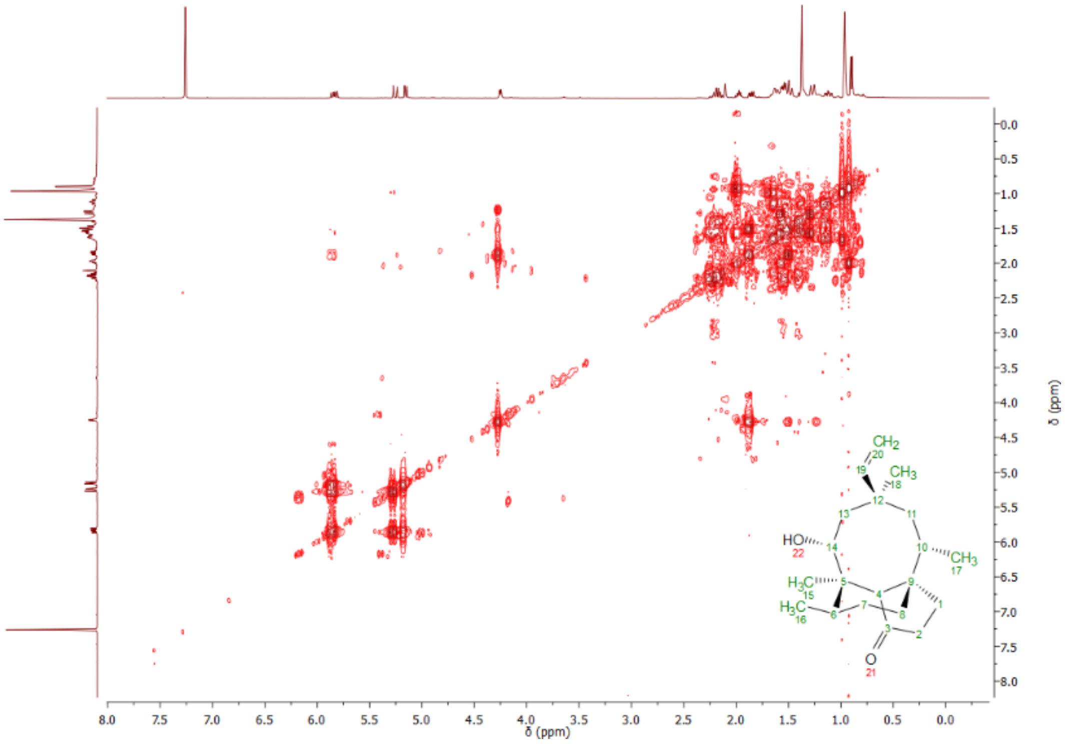

**Supplementary Figure 10.** COSY spectrum of **8** in $CDCl_3$ (500 MHz).

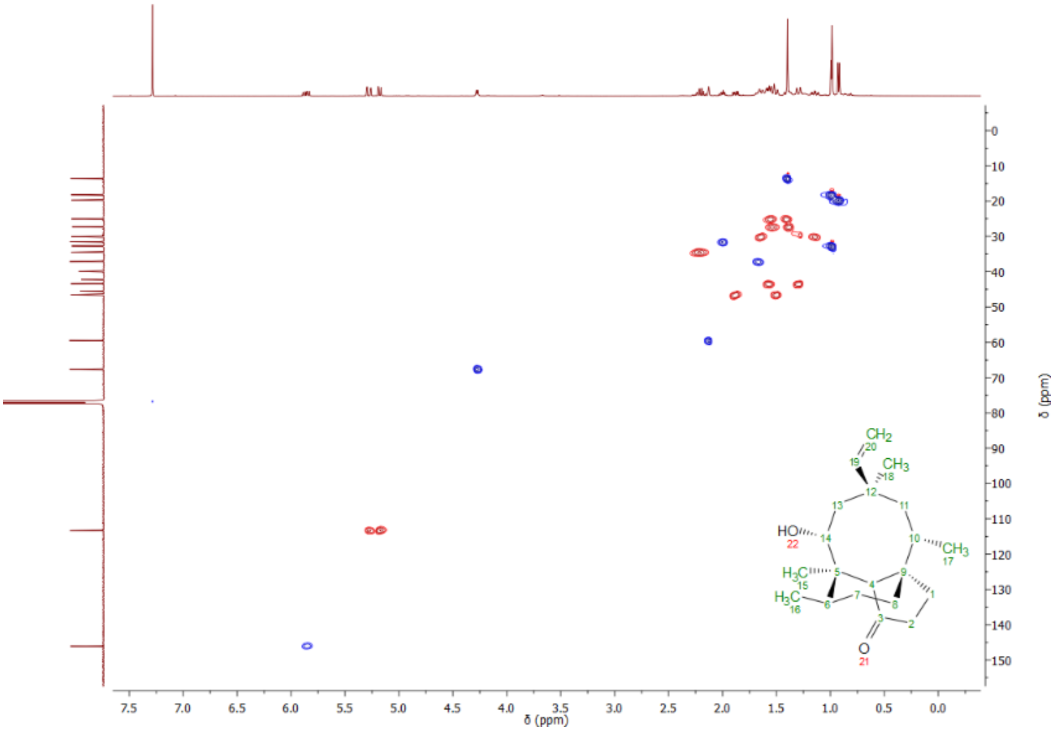

**Supplementary Figure 11.** HSQC spectrum of **8** in $CDCl_3$ (500 MHz).

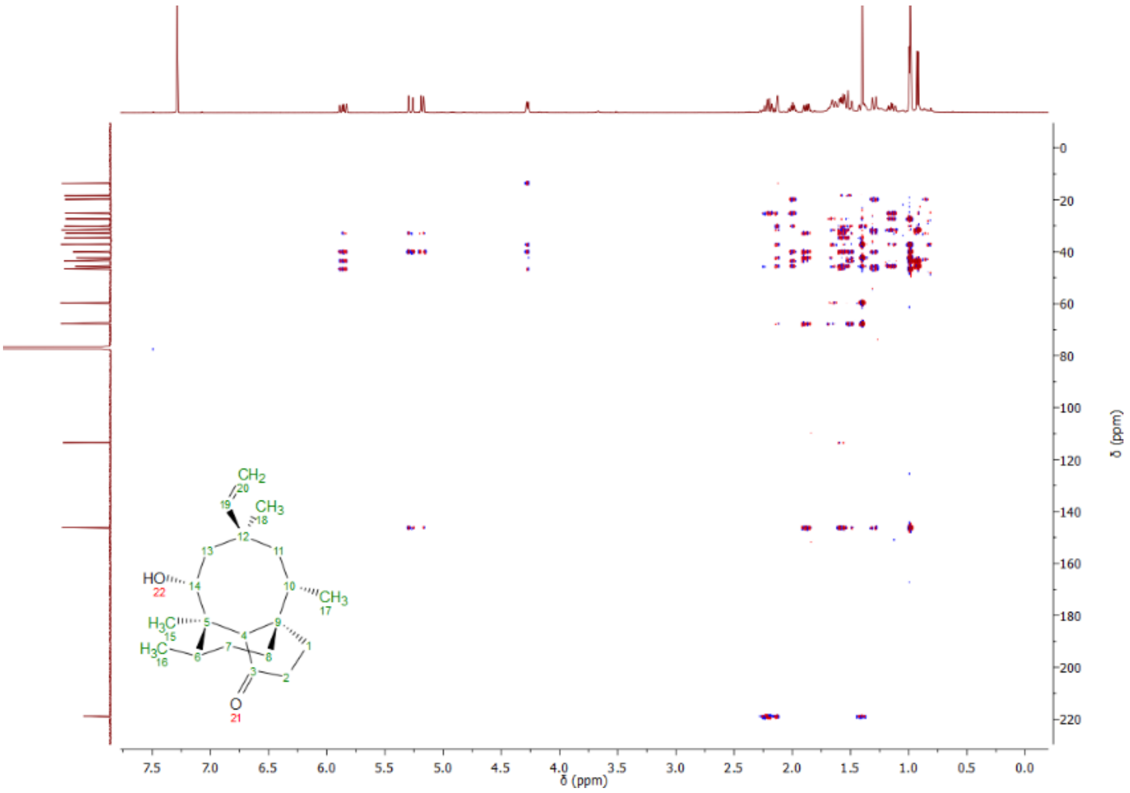

**Supplementary Figure 12.** HMBC spectrum of **8** in $CDCl_3$ (500 MHz).

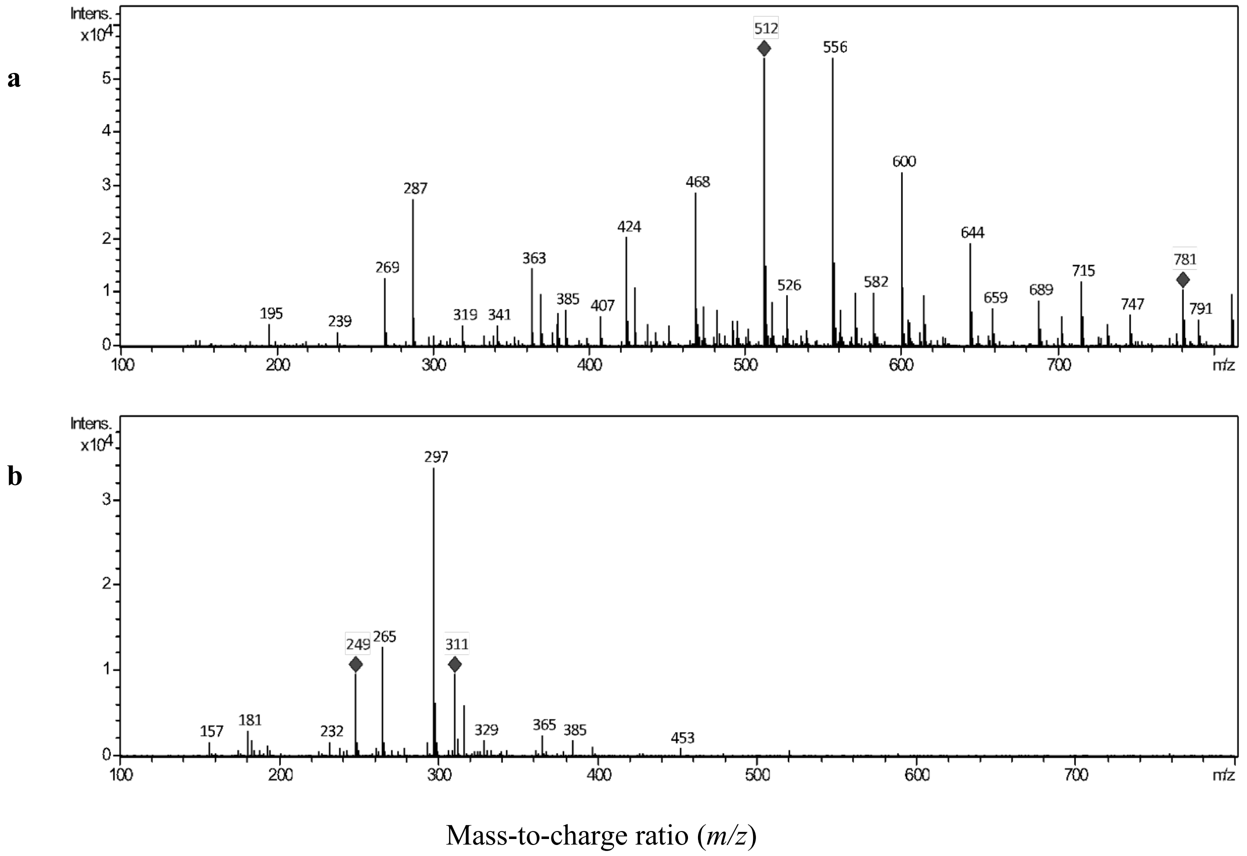

**Supplementary Figure 13.** Mass spectra in positive (a) and negative (b) ion mode of **9**.

**NMR data assignment of 9 in $CDCl_3$**

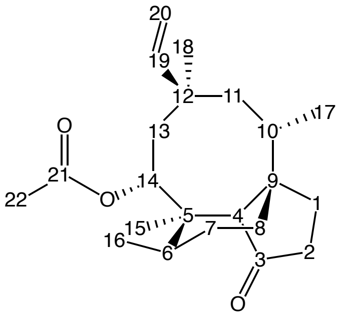

$\delta_H$ (500 MHz, $CDCl_3$) 6.21 (1H, dd, *J* 11.0, 17.6, 19-H), 5.57 (1H, d, *J* 8.0, 14-H), 5.17 (1H, d, *J* 11.0, 20-*H*H), 5.08 (1H, d, *J* 17.6, 20-H*H*), 2.29-2.16 (4H, m, 2-$H_2$, 4-H, 10-H), 2.03-1.96 (4H, m, 13-*H*H, 22-$H_3$), 1.71-1.54 (5H, m, 1-*H*H, 6-H, 7-*H*H, 8-*H*H, 11-*H*H), 1.48 (3H, s, 15-$H_3$), 1.41 (1H, m, 1-H*H*), 1.38-1.31 (2H, m, 7-H*H*, 11-H*H*), 1.27 (1H, m, 13-H*H*), 1.19-1.08 (1H, m, 8-H*H*), 1.00 (3H, s, 18-$H_3$), 0.93 (3H, d, *J* 6.7, 17-$H_3$), 0.74 (3H, d, *J* 5.9, 16-$H_3$). $\delta_C$ (125 MHz, $CDCl_3$) 218.6 (C-3), 169.7 (C-21), 146.1 (C-19), 112.6 (C-20), 69.2 (C-14), 59.0 (C-4), 47.0 (C-13), 45.7 (C-9), 42.9 (C-11), 41.9 (C-12), 38.9 (C-5), 37.1 (C-6), 34.7 (C-2), 32.2 (C-18), 31.1 (C-10), 30.1 (C-8), 27.0 (C-7), 25.1 (C-1), 22.1 (C-22), 19.6 (C-16), 15.9 (C-17), 14.5 (C-15). HRMS (ESI) calc. for $C_{22}H_{35}O_3^+$: 347.2586. Found 347.2581.

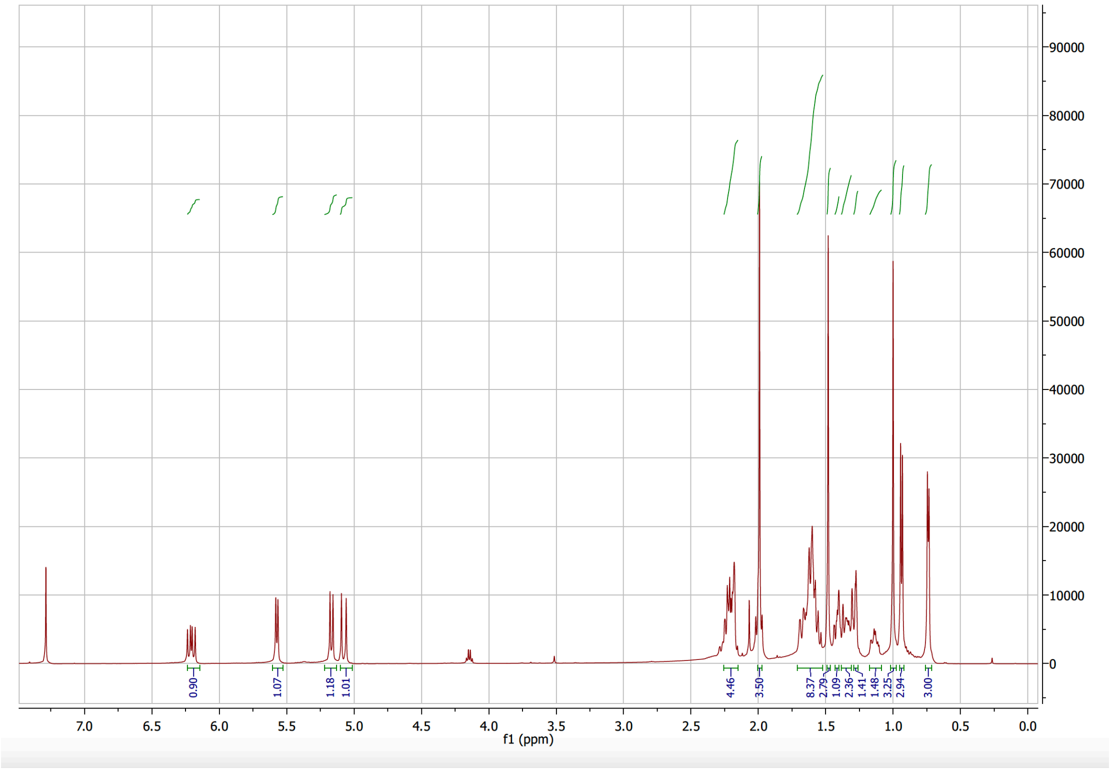

**Supplementary Figure 14.** [1]H-NMR spectrum of **9** in $CDCl_3$ (500 MHz).

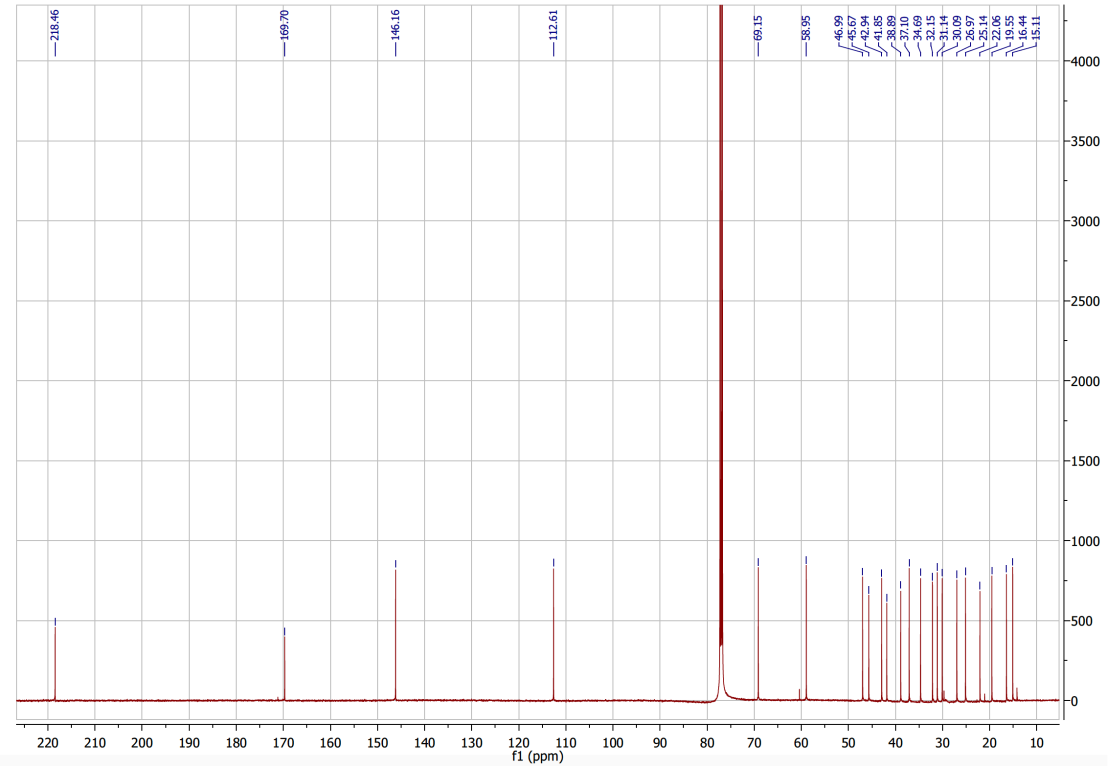

**Supplementary Figure 15.** [13]C-NMR spectrum of **9** in $CDCl_3$ (125 MHz).

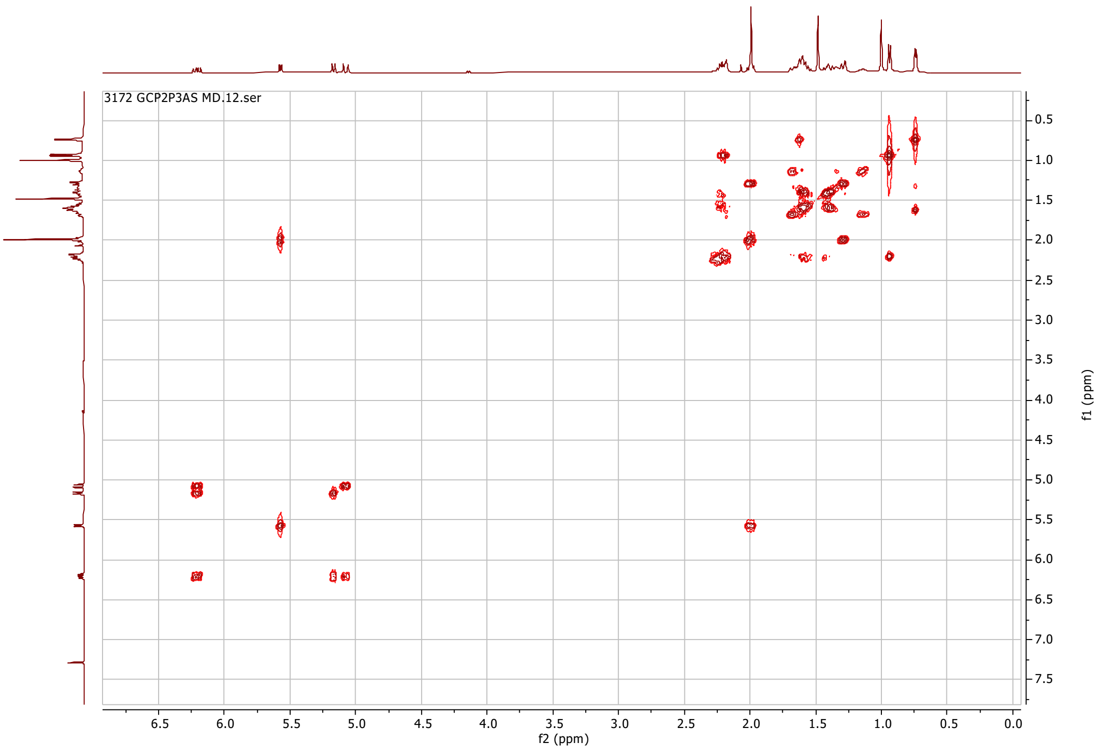

**Supplementary Figure 16.** COSY spectrum of **9** in $CDCl_3$ (500 MHz).

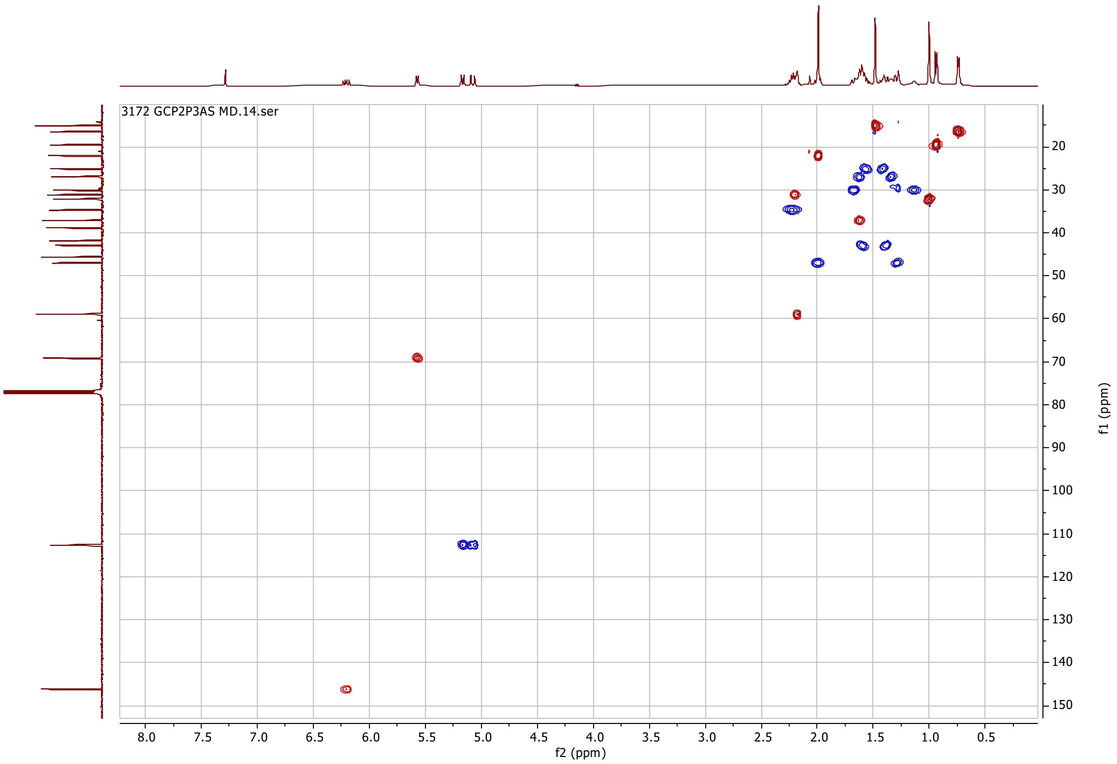

**Supplementary Figure 17.** HSQC spectrum of **9** in $CDCl_3$ (500 MHz).

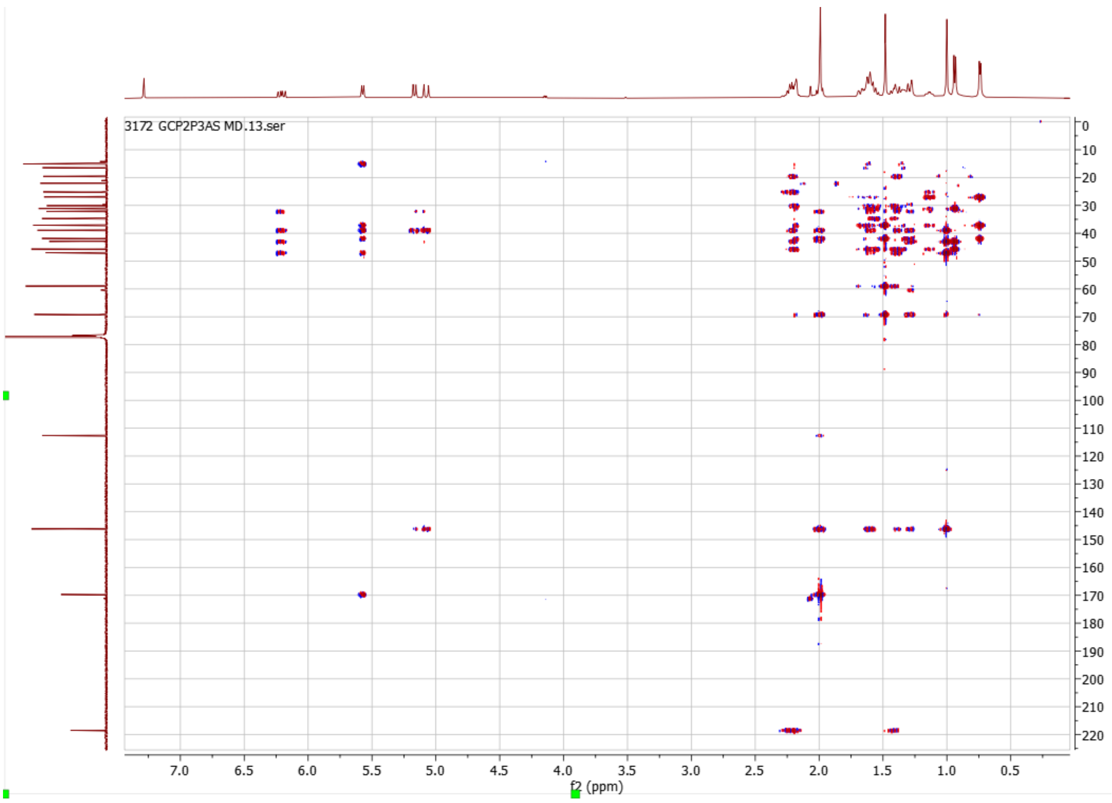

**Supplementary Figure 18.** HMBC spectrum of **9** in $CDCl_3$ (500 MHz).

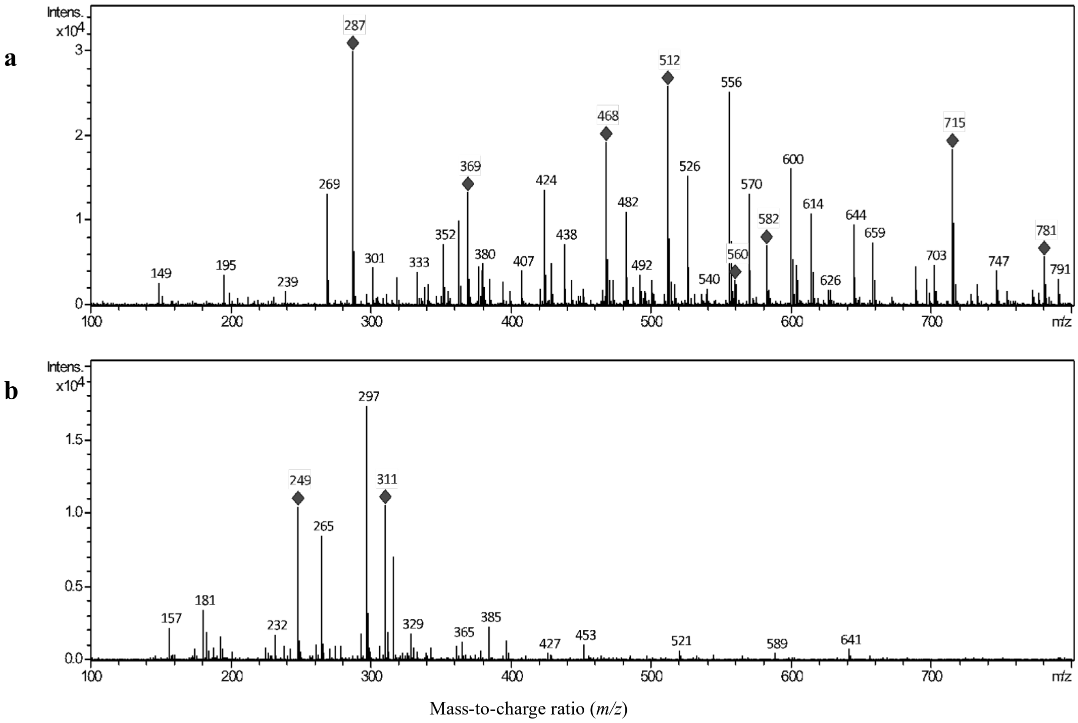

**Supplementary Figure 19.** Mass spectra in positive (a) and negative (b) ion mode of **10**.

**NMR data assignment of 10 in $CDCl_3$**

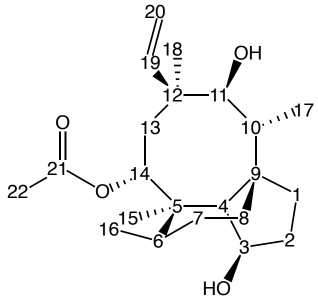

$\delta_H$ (500 MHz, $CDCl_3$) 6.57 (1H, dd, *J* 11.0, 17.4, 19-H), 5.57 (1H, d, *J* 9.4, 14-H), 5.34 (1H, dd, *J* 1.7, 11.0, 20-*H*H), 5.18 (1H, dd, *J* 1.7, 17.4, 20-*H*H), 4.56 (1H, t, *J* 5.35, 3-H), 3.18 (1H, s, 11-H), 2.27-2.15 (2H, m, 6-H, 10-H), 2.10-2.03 (1H, m, 13-*H*H), 1.98 (3H, s, 22-$H_3$), 1.96-1.81 (2H, m, 2-*H*H, 8-*H*H), 1.76-1.70 (1H, m, 2-H*H*), 1.69-1.61 (2H, m, 8-H*H,* 1-*H*H), 1.54-1.34 (4H, m, 4-H, 7-*H*H, 1-H*H,* 7-H*H*), 1.26 (1H, s, 13-H*H*), 1.23 (3H, s, 16-$H_3$), 1.16 (3H, s 18-$H_3$), 0.81 (3H, d, *J* 7.1, 17-$H_3$), 0.73 (3H, d, *J* 7.2,15-$H_3$). $\delta_C$ (125 MHz, $CDCl_3$) 170.0 (C-21), 139.7 (C-19), 116.5 (C-20), 77.3 (C-3), 74.9 (C-11), 69.8 (C-14), 51.0 (C-4), 46.0 (C-9), 45.4 (C-13), 44.9 (C-12), 41.1 (C-5), 36.6 (C-6), 35.6 (C-10), 34.3 (C-8), 32.7 (C-2), 31.8 (C-1), 27.7 (C-7), 26.1 (C-18), 21.9 (C-22), 17.6 (C-16), 16.9 (C-15), 12.2 (C-17). HRMS (ESI) calc. for $C_{22}H_{37}O_4^+$ 365.2692. Found 365.2684.

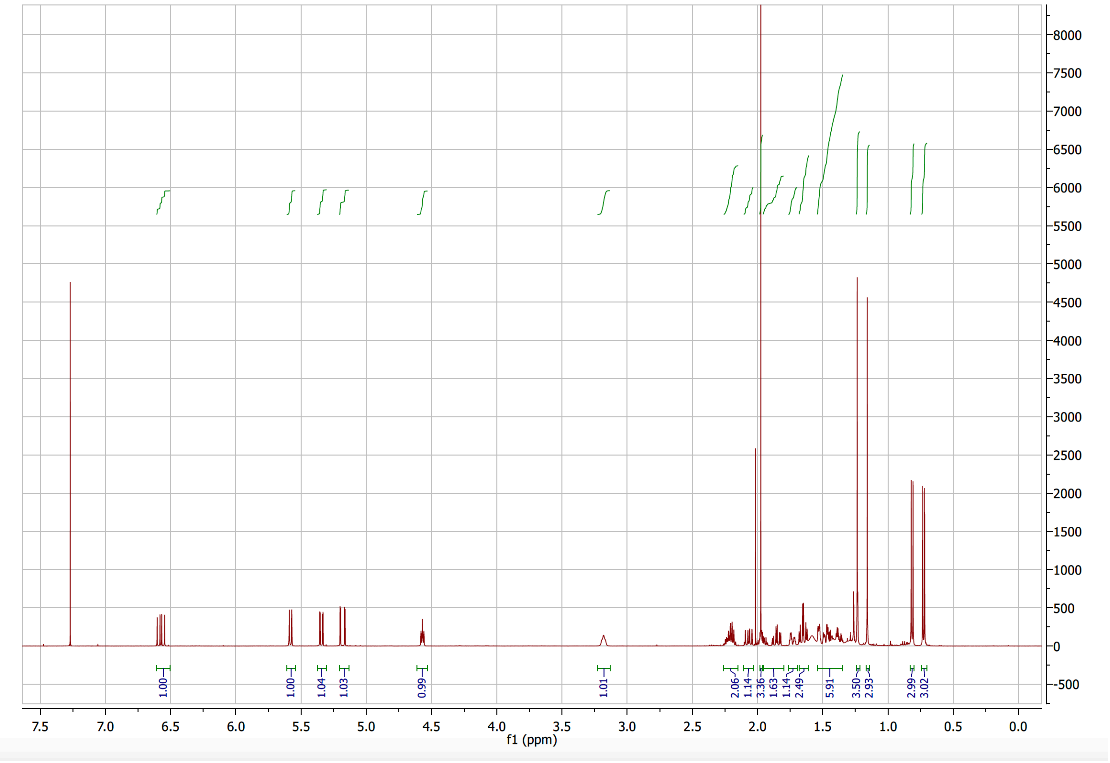

**Supplementary Figure 20.** $^{1}$H-NMR spectrum of **10** in $CDCl_3$ (500 MHz).

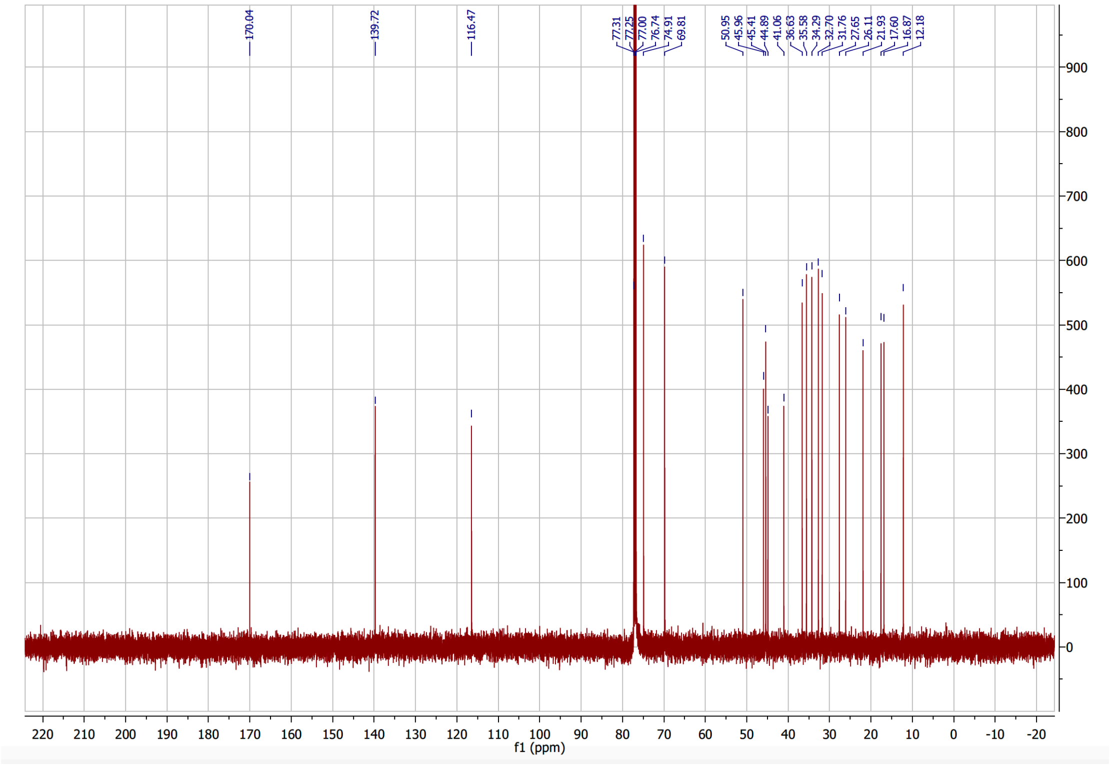

**Supplementary Figure 21.** $^{13}$C-NMR spectrum of **10** in $CDCl_3$ (125 MHz).

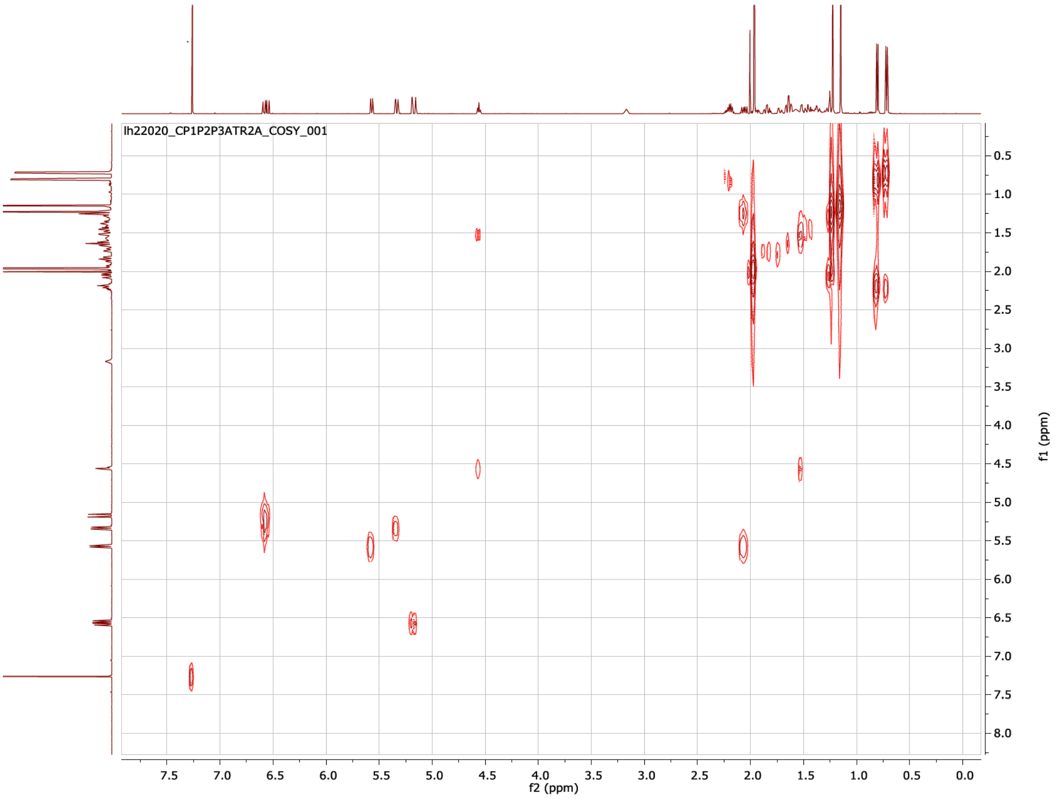

**Supplementary Figure 22.** COSY spectrum of **10** in $CDCl_3$ (500 MHz).

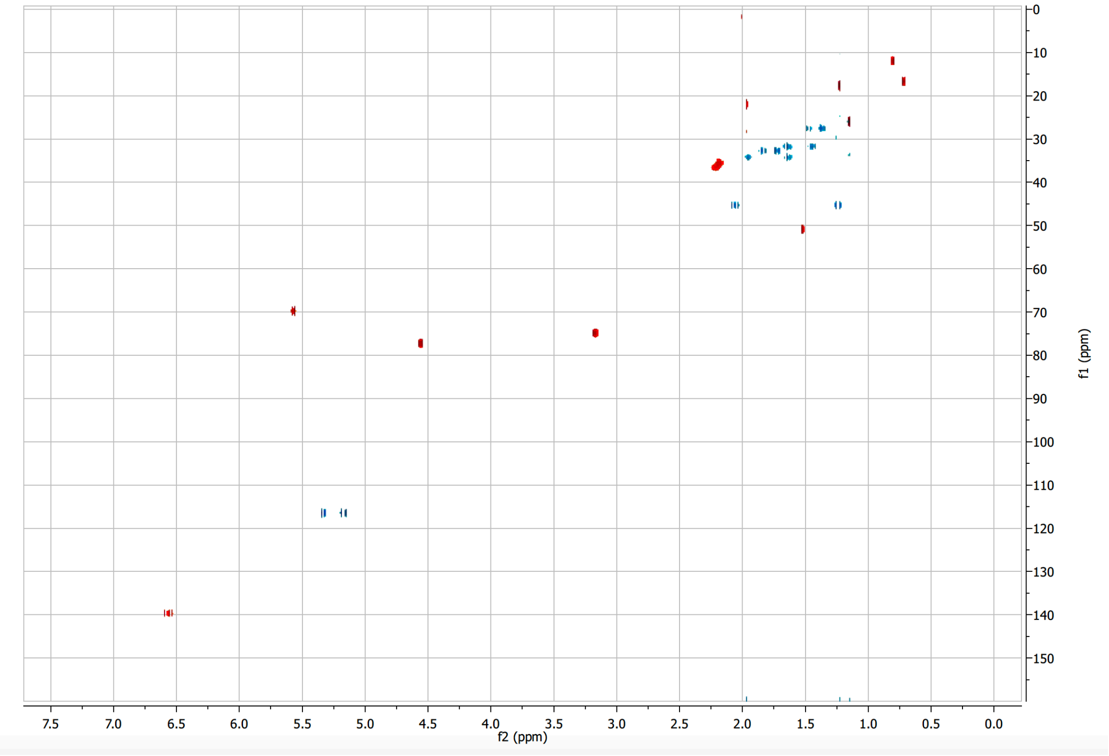

**Supplementary Figure 23.** HSQC spectrum of **10** in $CDCl_3$ (500 MHz).

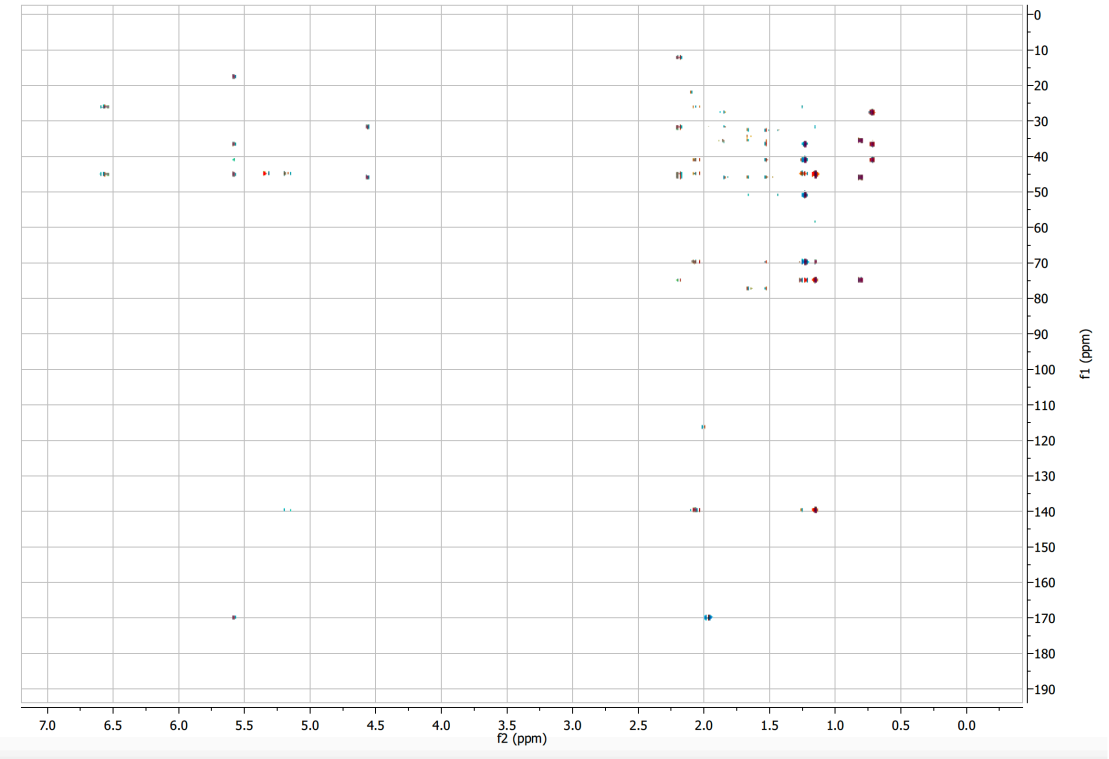

**Supplementary Figure 24.** HMBC spectrum of **10** in $CDCl_3$ (500 MHz).

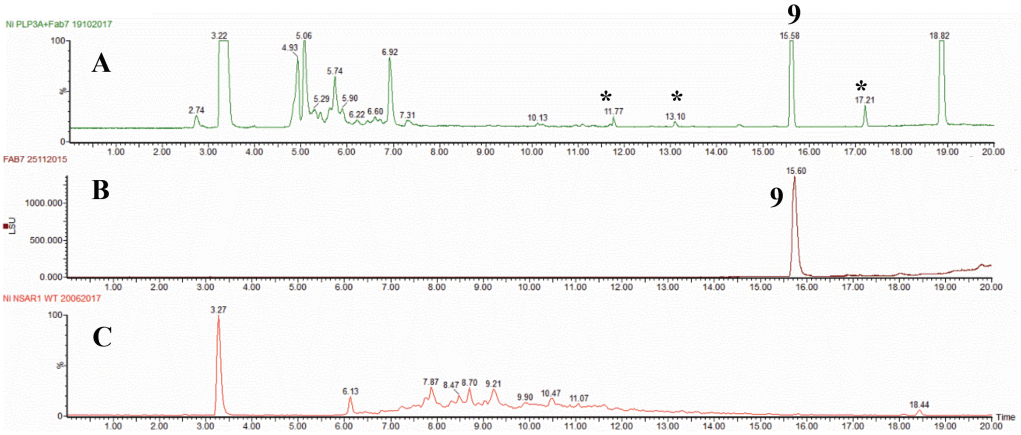

**Supplementary Figure 25.** ELSD chromatograms showing conversion of **8** to **9** in *A. oryzae* AP3. A) Feeding of *A. oryzae* AP3 with **8** gave successful production of acetylated **9**, but no 22-OH product was observed. Compounds with (*) were not related to pleuromutilin based on their m/z values; B) Purified metabolite **9**; C) *A. oryzae* NSAR1.

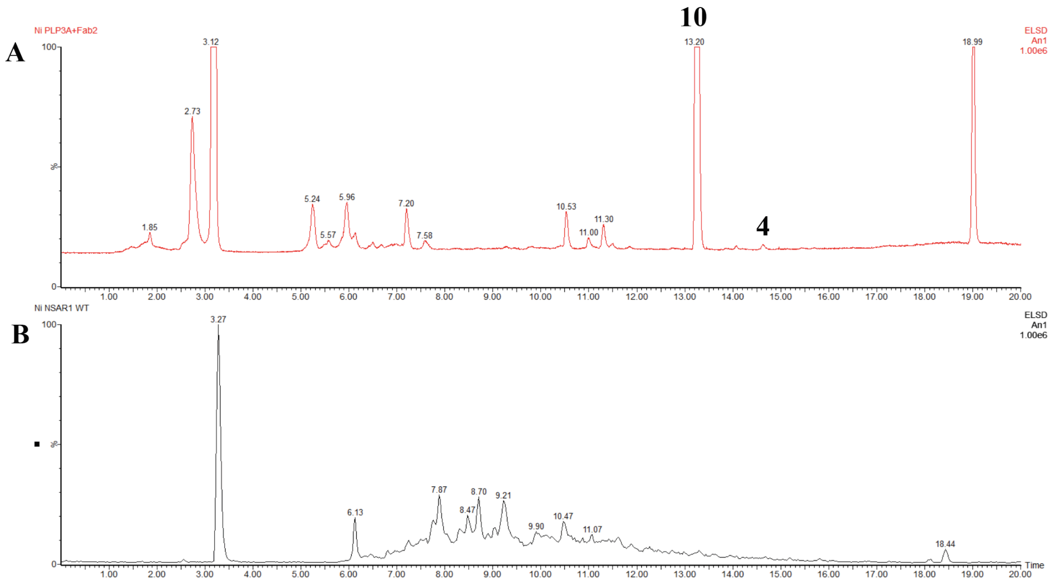

**Supplementary Figure 26.** ELSD chromatograms showing conversion of **4** to **10** in *A. oryzae* AP3. A) Feeding of *A. oryzae* AP3 with **4** gave successful production of acetylated **10**, but no 22-OH product was observed. B) *A. oryzae* NSAR1.

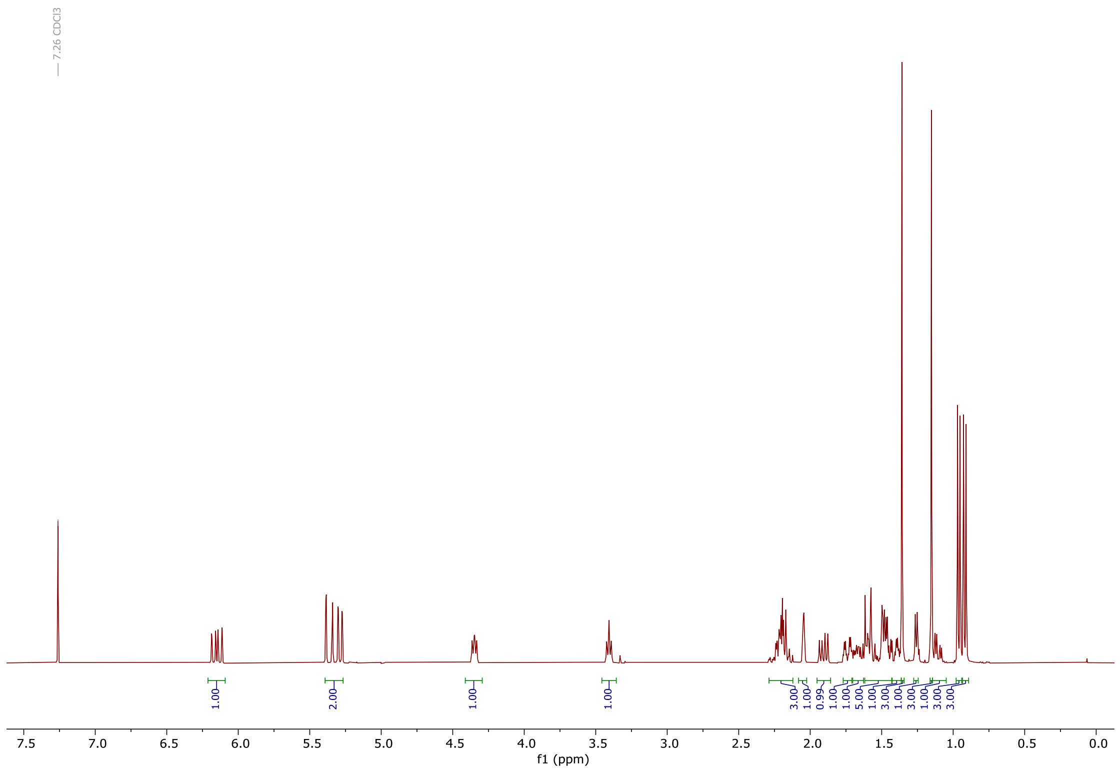

**Supplementary Figure 27. $^1$H-NMR spectrum of 5 in $CDCl_3$ (400 MHz).**

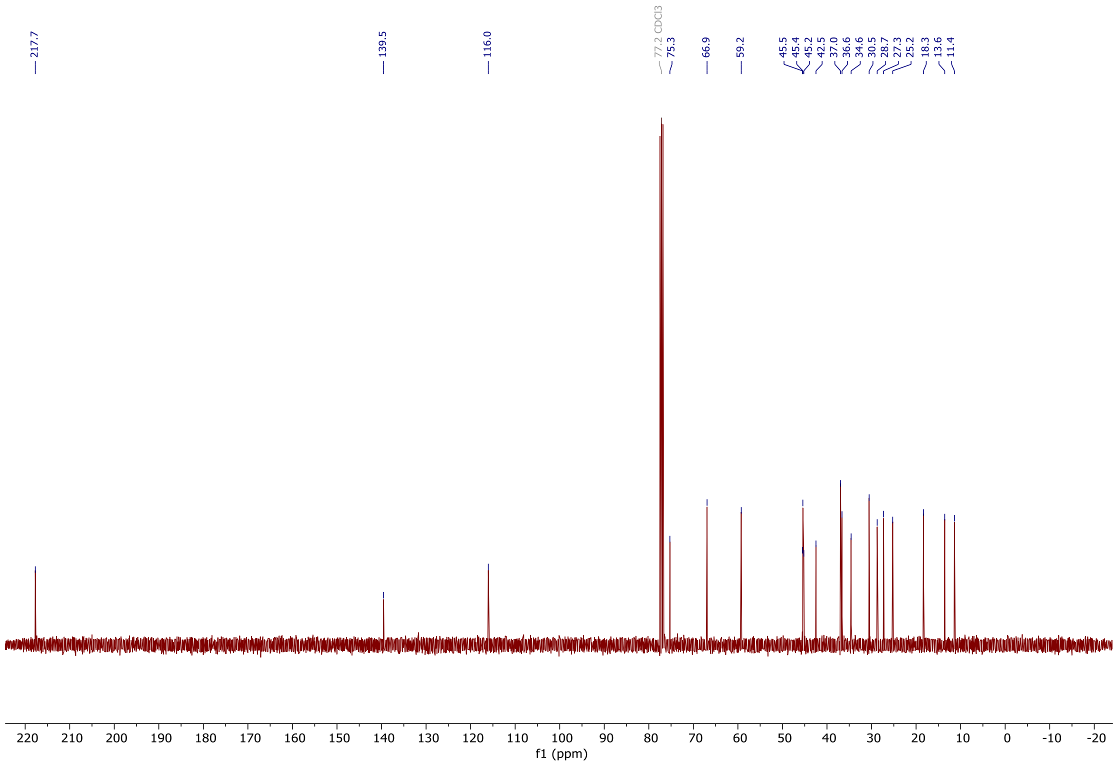

**Supplementary Figure 28. $^{13}$C-NMR spectrum of 5 in $CDCl_3$ (100 MHz).**

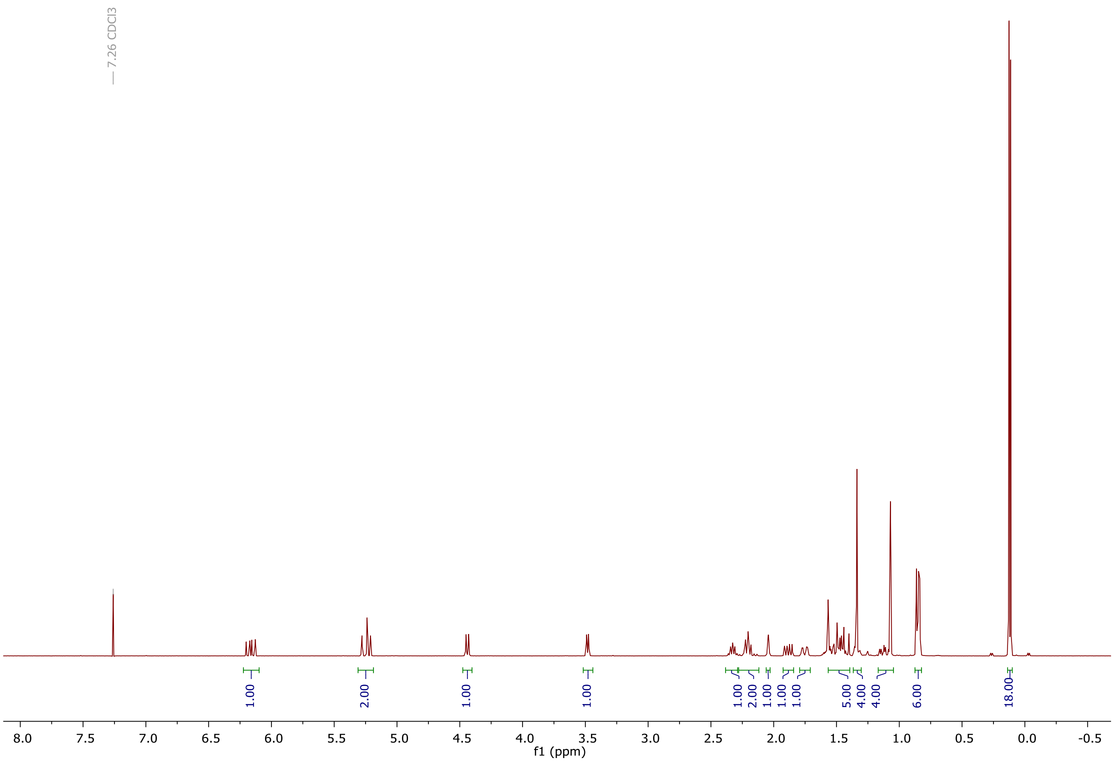

**Supplementary Figure 29. $^{1}$H-NMR spectrum of 12 in $CDCl_3$ (400 MHz).**

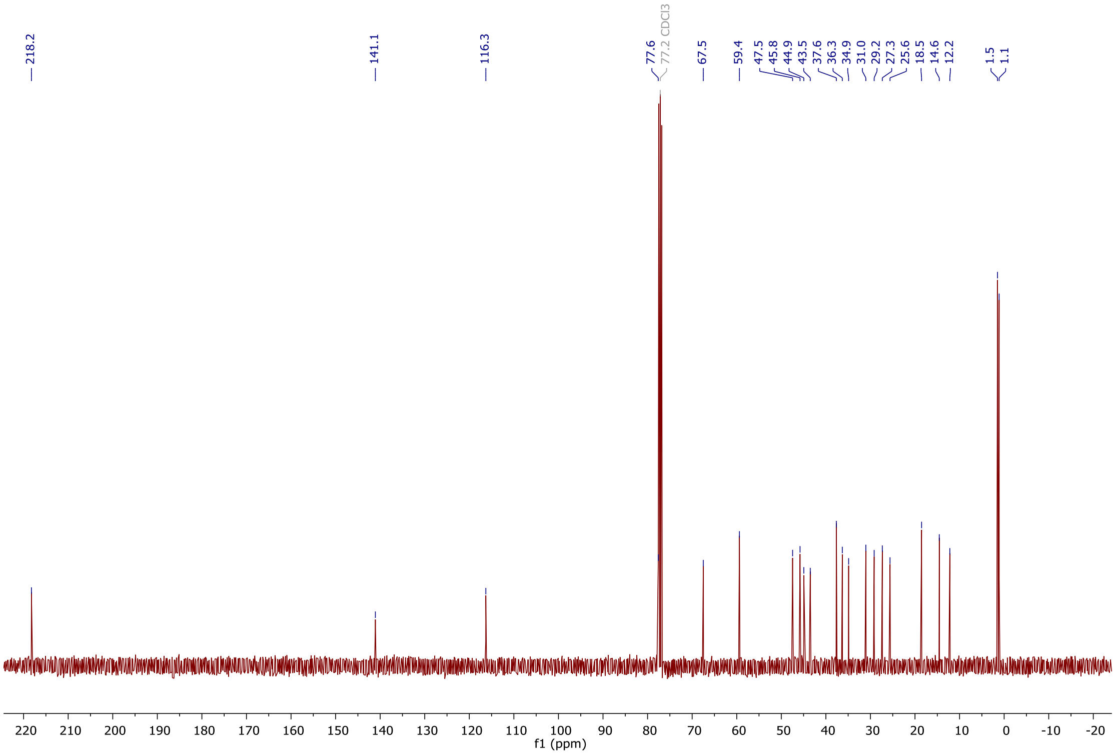

**Supplementary Figure 30. $^{13}$C-NMR spectrum of 12 in $CDCl_3$ (100 MHz).**

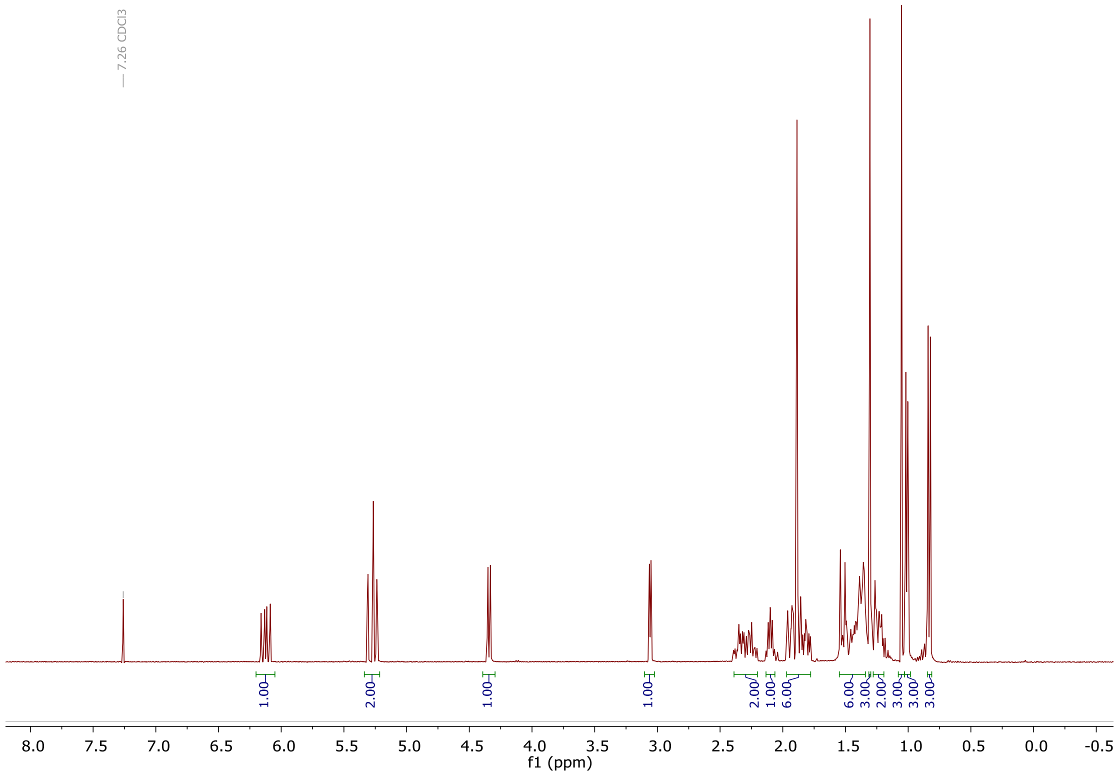

**Supplementary Figure 31. $^{1}$H-NMR spectrum of 14 in $CDCl_3$ (400 MHz).**

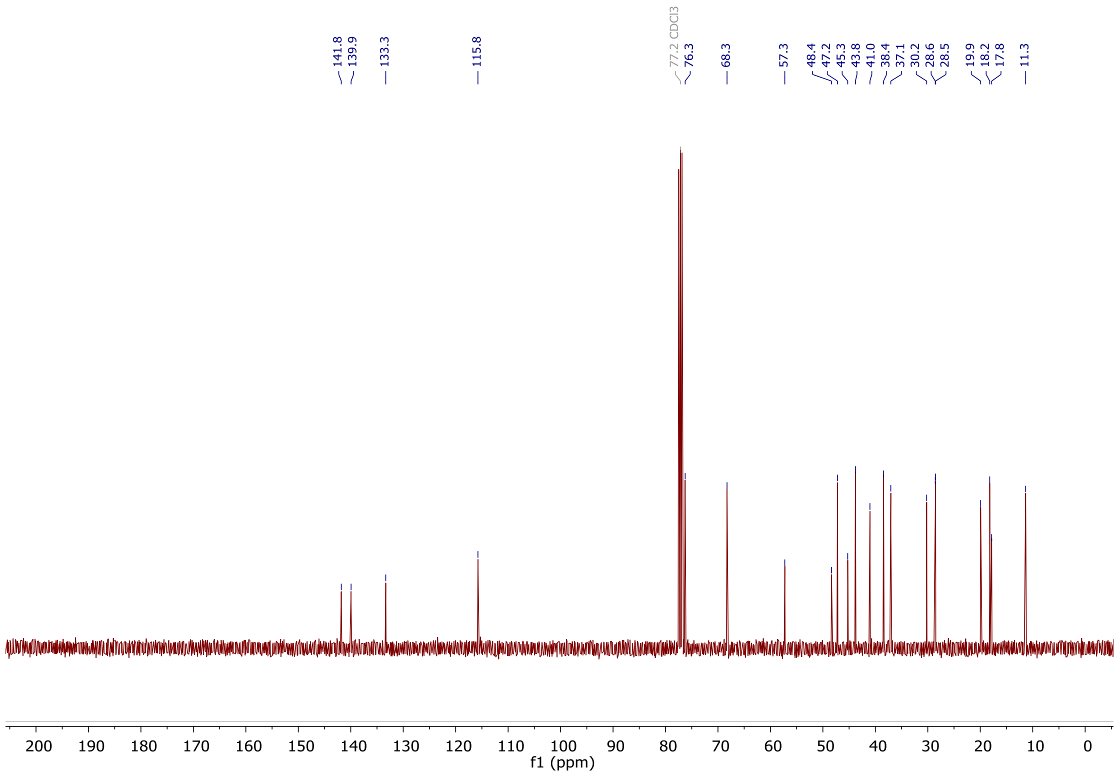

**Supplementary Figure 32. $^{13}$C-NMR spectrum of 14 in $CDCl_3$ (100 MHz).**

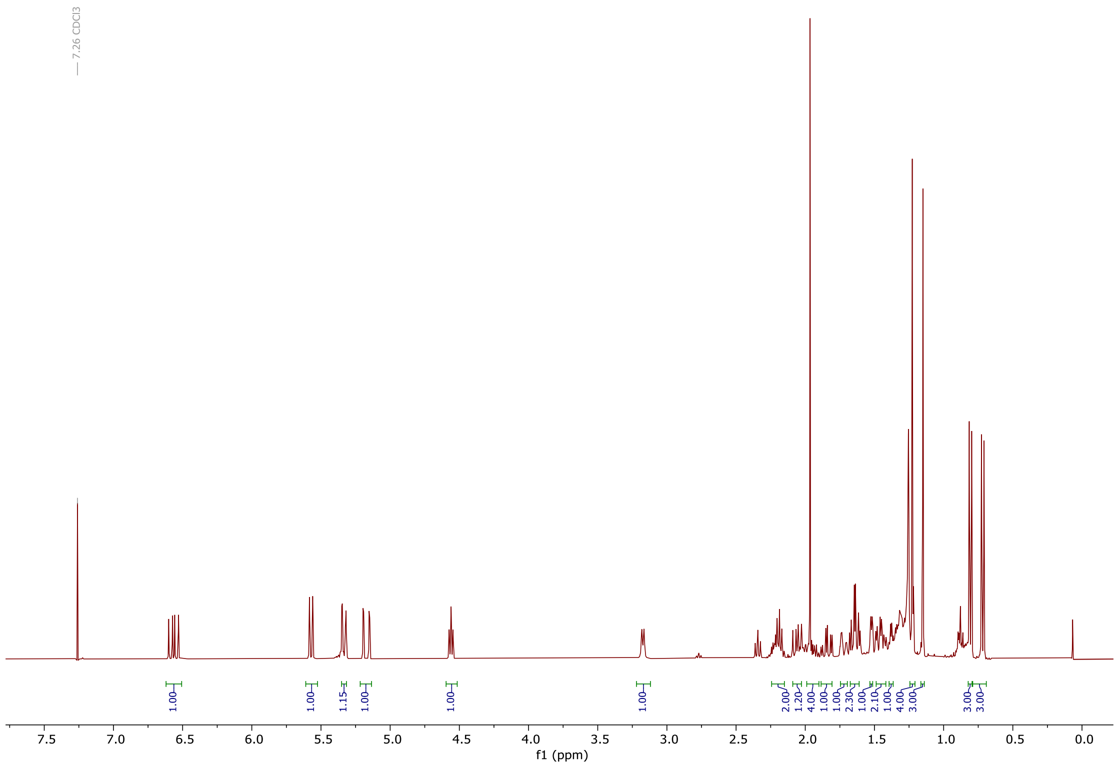

**Supplementary Figure 33.** **$^{1}$H-NMR spectrum of 15 in $CDCl_3$ (400 MHz).**

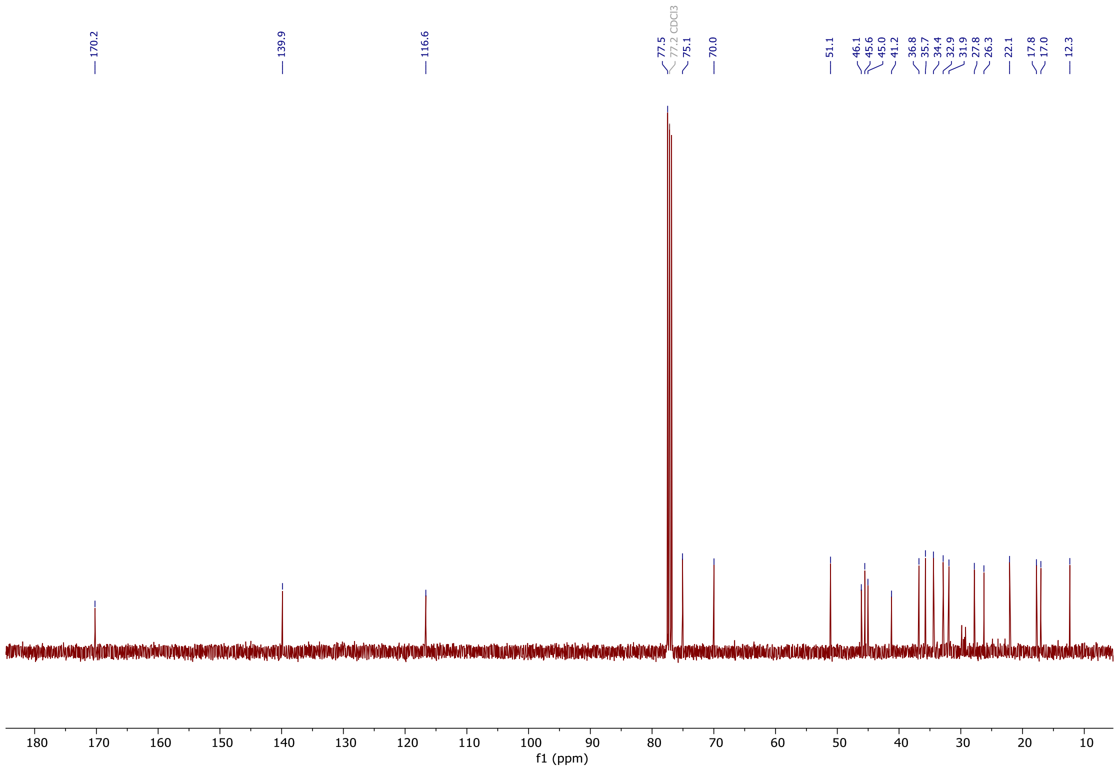

**Supplementary Figure 34.** **$^{13}$C-NMR spectrum of 15 in $CDCl_3$ (100 MHz).**

### 2. Supplementary Methods

**Synthesis of mutilin 5**

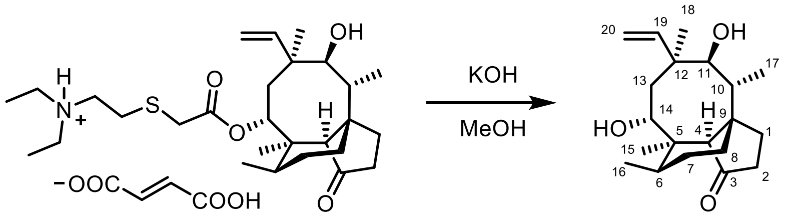

tiamulin hydrogen fumarate mutilin **5**

Potassium hydroxide (7.50 g, 134 mmol) was added to a solution of tiamulin hydrogen fumarate (3.45 g, 5.65 mmol) in methanol (100 mL). The reaction was heated to reflux (65 °C) for 18 hours, before cooling to room temperature and pouring into water (50 mL). The resulting solution was extracted with dichloromethane (3 × 50 mL) and the combined organic layers were washed with a saturated aqueous solution of $NaHCO_3$ (50 mL), dried over $MgSO_4$, filtered, and concentrated *in vacuo.* The crude material was purified by column chromatography (20% EtOAc in petrol) to give mutilin **5** (1.22 g, 68%) as a white solid.

$\delta_H$ (400 MHz, $CDCl_3$) 6.15 (1H, ddd, *J* 18.0, 11.0, 0.5, 19-H), 5.36 (1H, dd, *J* 18.0, 1.5, 20-*H*H), 5.29 (1H, dd, *J* 11.0, 1.5, 20-H*H*), 4.35 (1H, dd, *J* 8.0, 6.0, 14-H), 3.41 (1H, dd, *J* 7.5, 5.5, 11-H), 2.29-2.10 (3H, m, 2-$H_2$, 10-H), 2.05 (1H, d, *J* 3.0, 4-H), 1.91 (1H, dd, *J* 16.0, 8.0, 13-*H*H), 1.74 (1H, dq, *J* 14.5, 3.0, 8-*H*H), 1.67 (1H, m, 6-H), 1.63-1.42 (5H, m, 1-$H_2$, 7-*H*H, 13-H*H*, O*H*), 1.39 (1H, m, 7-H*H*), 1.36 (3H, s, 15-$H_3$), 1.26 (1H, d, *J* 5.5, O*H*), 1.12 (1H, m, 8-H*H*), 1.15 (3H, s, 18-$H_3$), 0.96 (3H, d, *J* 7.0, 16-$H_3$), 0.92 (3H, d, *J* 7.0, 17-$H_3$). $\delta_C$ (101 MHz, $CDCl_3$) 217.7 (C-3), 139.5 (C-19), 116.0 (C-20), 75.3 (C-11), 66.9 (C-14), 59.2 (C-4), 45.5 (C-9), 45.4 (C-12), 45.2 (C-13), 42.5 (C-5), 37.0 (C-6), 36.6 (C-10), 34.6 (C-2), 30.5 (C-8), 28.7 (C-18), 27.3 (C-7), 25.2 (C-1), 18.3 (C-16), 13.6 (C-15), 11.4 (C-17). HRMS (ESI) calc. for $C_{20}H_{32}O_3Na$ $[M+Na]^+$ 343.2244. Found 343.2245. M.p. 190-191 °C (from EtOH) [Lit. 192 °C].[153] $[\alpha]_D^{21}$ = +31.0 (*c* 1.00, $CHCl_3$) [Lit. +69 (*c* 0.07, $CHCl_3$)].[242] IR ($\nu_{max}/cm^{-1}$) (neat): 3471 (alcohol O–H), 2929 (alkane C–H), 1727 (ketone C=O).

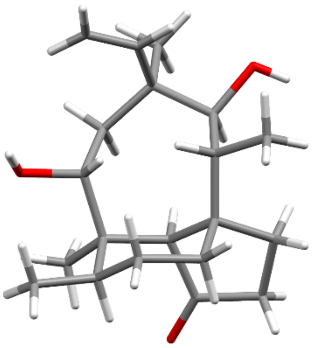

Spectroscopic data in accordance with the literature data.[1]

X-ray crystal structure obtained following recrystallisation from dichloromethane.

Space Group: $P2_12_12_1$ (orthorhombic).

**Synthesis of TMS-mutilin 12**

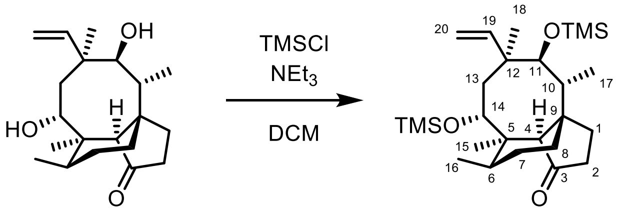

mutilin **5** TMS-mutilin **12**

Mutilin **5** (500 mg, 1.56 mmol) and Trimethylsilyl chloride (0.99 mL, 7.80 mmol) were dissolved in dichloromethane (10 mL). Triethylamine (1.08 mL, 7.80 mmol) was added dropwise and the solution stirred for 16 hours at room temperature. A saturated aqueous solution of $NaHCO_3$ (10 mL) was added to the reaction mixture and the layers were separated. The aqueous phase was extracted with dichloromethane (3 × 10 mL) and the combined organic layers were dried over $MgSO_4$, filtered, and concentrated *in vacuo*. The crude material was purified by column chromatography (5% EtOAc in petrol) to give TMS-mutilin **12** (668 mg, 92%) as a colourless oil.

$\delta_H$ (400 MHz, $CDCl_3$) 6.16 (1H, dd, *J* 17.5, 11.0, 19-H), 5.26 (1H, dd, *J* 17.5, 1.5, 20-*H*H), 5.22 (1H, dd, *J* 11.0, 1.5, 20-H*H*), 4.44 (1H, d, *J* 8.0, 14-H), 3.48 (1H, d, *J* 6.0, 11-H), 2.33 (1H, app. pent, *J* 7.0, 10-H), 2.25-2.12 (2H, m, 2-$H_2$), 2.04 (1H, d, *J* 3.0, 4-H), 1.88 (1H, dd, *J* 16.0, 8.0, 13-*H*H), 1.75 (1H, dq, *J* 14.5, 3.0, 8-*H*H), 1.57 (1H, m, 6-H), 1.55-1.39 (2H, m, 1-$H_2$, 7-*H*H), 1.42 (1H, d, *J* 16.0, 13-H*H*), 1.35 (1H, m, 7-H*H*), 1.34 (3H, s, 15-$H_3$), 1.12 (1H, td, *J* 14.0, 4.5, 8-H*H*), 1.07 (3H, s, 18-$H_3$), 0.85 (3H, d, *J* 7.0, 16-$H_3$), 0.85 (3H, d, *J* 7.0, 17-$H_3$), 0.13 (9H, s, $(SiCH_3)_3$), 0.11 (9H, s, $(SiCH_3)_3$). $\delta_C$ (101 MHz, $CDCl_3$) 218.2 (C-3), 141.1 (C-19), 116.3 (C-20), 77.6 (C-11), 67.5 (C-14), 59.4 (C-4), 47.5 (C-13), 45.8 (C-9), 44.9 (C-12), 43.5 (C-5), 37.6 (C-6), 36.5 (C-10), 34.9 (C-2), 31.0 (C-8), 29.2 (C-18), 27.3 (C-7), 25.6 (C-1), 18.5 (C-16), 14.6 (C-15), 12.2 (C-17), 1.5 and 1.1 ($SiCH_3$). HRMS (ESI) calc. for $C_{26}H_{48}O_2Si_2Na$ $[M+Na]^+$ 487.3034. Found 487.3045. $[\alpha]_D^{21}$ = +44.0 (*c* 0.75, $CHCl_3$). IR ($\nu_{max}/cm^{-1}$) (neat): 2956 (alkane C–H), 1736 (ketone C=O).

$^1$H NMR data in accord with the literature data.[2]

**Synthesis of alkene 14**

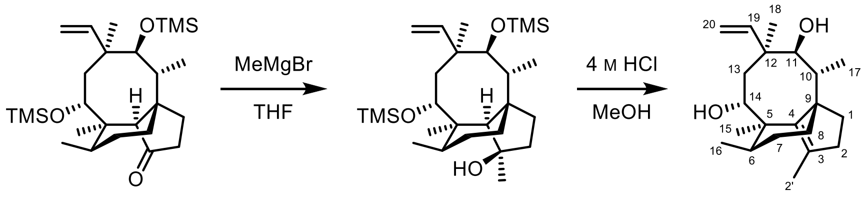

TMS-mutilin **12** (400 mg, 0.861 mmol) was dissolved in tetrahydrofuran (10 mL) and cooled to 0 °C. A solution of methylmagnesium bromide in $Et_2O$ (3 M, 1.43 mL, 4.30 mmol) was added dropwise and the solution warmed to room temperature and stirred for 16 hours. The reaction was quenched with water (20 mL) and extracted with dichloromethane (3 × 20 mL). The combined organic layers were dried over $MgSO_4$, filtered, and concentrated *in vacuo*.

The resulting material was dissolved in a solution of HCl in MeOH (4 M, 7.5 mL) and stirred for 16 hours at room temperature. The reaction mixture was neutralised with aqueous NaOH solution and extracted with dichloromethane (3 × 15 mL). The combined organic layers were dried over $MgSO_4$, filtered, and concentrated *in vacuo*. The crude material was purified by column chromatography (20% EtOAc in petrol) to give alkene **14** (153 mg, 56%) as a white solid.

**Alkene 14**

$\delta_H$ (400 MHz, $CDCl_3$) 6.12 (1H, dd, *J* 18.0, 11.0, 19-H), 5.29 (1H, dd, *J* 18.0, 1.5, 20-*H*H), 5.25 (1H, dd, *J* 11.0, 1.5, 20-H*H*), 4.34 (1H, d, *J* 8.0, 14-H), 3.06 (1H, d, *J* 5.5, 11-H), 2.35 (1H, ddd, *J* 17.0, 11.0, 4.5, 2-*H*H), 2.25 (1H, ddd, *J* 17.0, 10.0, 7.0, 2-H*H*), 2.10 (1H, qd, *J* 7.0, 5.5, 10-H), 1.98-1.77 (3H, m, 1-*H*H, 8-*H*H, 13-H*H*), 1.89 (3H, s, 2'-$H_3$), 1.56-1.32 (5H, m, 6-H, 7-$H_2$, 2 × O*H*), 1.52 (1H, d, *J* 15.0, 13-H*H*), 1.31 (3H, s, 15-$H_3$), 1.30-1.17 (2H, m, 1-H*H*, 8-H*H*), 1.05 (3H, s, 18-$H_3$), 1.01 (3H, d, *J* 6.5, 16-$H_3$), 0.83 (3H, d, *J* 6.5, 17-$H_3$). $\delta_C$ (101 MHz, $CDCl_3$) 141.8 (C-4), 139.9 (C-19), 133.3 (C-3), 115.8 (C-20), 76.3 (C-11), 68.3 (C-14), 57.3 (C-9), 48.4 (C-5), 47.2 (C-13), 45.3 (C-12), 43.8 (C-6), 41.0 (C-2), 38.4 (C-8), 37.1 (C-10), 30.2 (C-1), 28.6 and 28.5 (C-7 and C-18), 19.9 (C-15), 18.2 (C-16), 17.8 (C-2'), 11.3 (C-17). HRMS (ESI) calc. for $C_{21}H_{34}O_2Na$ $[M+Na]^+$ 341.2451. Found 341.2467. M.p. 118-120 °C (from $CHCl_3$). $[\alpha]_D^{22} = -55.0$ (*c* 1.00, $CHCl_3$). IR ($\nu_{max}/cm^{-1}$) (neat): 3466 (alcohol O–H), 2921 (alkane C–H).

**Supplementary Table 1.** Summary of expression vectors used in this study to express genes from C. *passeckerianus* in *A. oryzae*. Selectable marker *argB*: *A. nidulans* ornithine carbamoyltransferase (OCTase) gene; *adeA*: *A. oryzae* phosphoribosyl-aminoimidazole-succinocarboxamide synthetase gene; *bar*: *Streptomyces spp.* phosphinothricin acetyl-transferase gene. Promoter Padh: *A. oryzae* alcohol dehydrogenase; PgpdA: *A. nidulans* glyceraldehyse 3'-phosphate dehydrogenase; Peno: *A. oryzae* enolase.

| **Backbone vector** | **Selectable marker gene** | **Promoter** | **Gene** | **Plasmid name** |
|---|---|---|---|---|
| pTYGSarg[3] | *argB* | Padh<br>Peno | *Pl-ggs*<br>*Pl-cyc* | pTYGSargGC[4] |
| pTYGSade[3] | *adeA* | Padh<br>PgpdA<br>Peno | *Pl-p450-1*<br>*Pl-p450-2*<br>*Pl-p450-3* | pTYGSadeP1P2P3[4] |
| | | Padh<br>Peno | *Pl-p450-2*<br>*Pl-p450-3* | pTYGSadeP2P3 |
| | | Padh | *Pl-p450-2* | pTYGSadeP2 |
| | | Padh | *Pl-p450-3* | pTYGSadeP3[4] |
| pTYGSbar[3] | *bar* | Padh<br>Peno | *Pl-atf*<br>*Pl-sdr* | pTYGSbarAS[4] |
| | | Padh | *Pl-atf* | pTYGSbarA[4] |
| | | Padh | *Pl-sdr* | pTYGSbarS[4] |

**Supplementary Table 2.** List of primers used in this study. Nucleotides in italics represent sequences used to generate overlapping regions with plasmid backbones for yeast-based homologous recombination.

| **Primer** | **DNA Sequence (5’->3’)** | **Description** |
|---|---|---|
| Pl-ggs FF | ATGAGAATACCTAACGTCTTTCTCT | Screening primers for the amplification of Pl-ggs |
| Pl-ggs RR | CTA CTC TGC GAT GTA CAA CTT TTC C | |
| Pl-cyc FF | ATG GGT CTA TCT GAA GAT CTT CAT G | Screening primers for the amplification of Pl-cyc |
| Pl-cyc RR | TCA ATG GTG GAT TCC ATT GCT CCC G | |
| Pl-p450-1 FF | ATG CTG TCC GTC GAC CTC CCG TCT G | Screening primers for the amplification of Pl-p450-1 |
| Pl-p450-1 RR | CTA CAA CGC AGC GAA CGC TTC CTT A | |
| Pl-p450-2 FF | ATG AAT CTT TCT GCT CTG AAG GCT G | Screening primers for the amplification of Pl-p450-2 |
| Pl-p450-2 RR | CTA ATA GTC TGC AAC ATC GTG GAT C | |
| Pl-p450-3 FF | ATG GCT CCG TCA ACG GAA CGT GCT C | Screening primers for the amplification of Pl-p450-3 |
| Pl-p450-3 RR | CTA GCC ACT AGC AGG CTT CGT GAA C | |
| Pl-atf FF | ATG AAG CCC TTC TCA CCA GAA CTT C | Screening primers for the amplification of Pl-atf |
| Pl-atf RR | CTA CTG TGC TAC ACG AGG GGG ATT C | |
| Pl-sdr FF | ATG GAA GGC AAG GTC GCA ATC GTC A | Screening primers for the amplification of Pl-sdr |
| Pl-sdr RR | CTA AAT GAC ACT CCA CCC GTT ATC G | |
| Padh-Pl-p450-2 FF | *TTTCTTTCAACACAAGATCCCAAAGTCAAA*ATGAATCTTTCTGCTCTGAA | Amplification of Pl-p450-2 for assembly of pTYGSadeP2P3 |
| Pl-p450-2-TgpdA RR | *ACGACAATGTCCATATCATCAATCATGACC*CTAATAGTCTGCAACATCGT | |
| Peno- Pl-p450-3 FF | *GTCGACTGACCAATTCCGCAGCTCGTCAAA*ATGGCTCCGTCAACGGAACG | Amplification of Pl-p450-2 for assembly of pTYGSadeP2P3 |
| Pl-p450-3-Teno RR | *GGTTGGCTGGTAGACGTCATATAATCATAC*CTAGCCACTAGCAGGCTTCG | |
| Padh-Pl-p450-2 FF | *TTTCTTTCAACACAAGATCCCAAAGTCAAA*ATGAATCTTTCTGCTCTGAA | Amplification of Pl-p450-2 for assembly of pTYGSadeP2 |
| Pl-p450-2-Teno RR | *GGTTGGCTGGTAGACGTCATATAATCATAC*CTAATAGTCTGCAACATCGT | |

**Supplementary Table 3.** List of *A. oryzae* strains generated in this study and relative expression vectors used to introduce the relevant pleuromutilin biosynthetic genes.

| *A. oryzae* strain | Heterologous genes from *C. passeckerianus* | Plasmids used for heterologous expression |
|---|---|---|
| GCP2 | *Pl-ggs, Pl-cyc, Pl-p450-2* | pTYGSargGC, pTYGSadeP2 |
| GCP2S | *Pl-ggs, Pl-cyc, Pl-p450-2, Pl-sdr* | pTYGSargGC, pTYGSadeP2, pTYGSbarS |
| GCP2P3SA | *Pl-ggs, Pl-cyc, Pl-p450-2, Pl-p450-3, Pl-sdr, Pl-atf* | pTYGSargGC, pTYGSadeP2P3, pTYGSbarAS |
| GCP1P2P3A | *Pl-ggs, Pl-cyc, Pl-p450-1, Pl-p450-2, Pl-p450-3, Pl-atf* | pTYGSargGC, pTYGSadeP1P2P3, pTYGSbarA |
| AP3 | *Pl-atf*, *Pl-p450-3* | pTYGSbarA, pTYGSadeP3 |

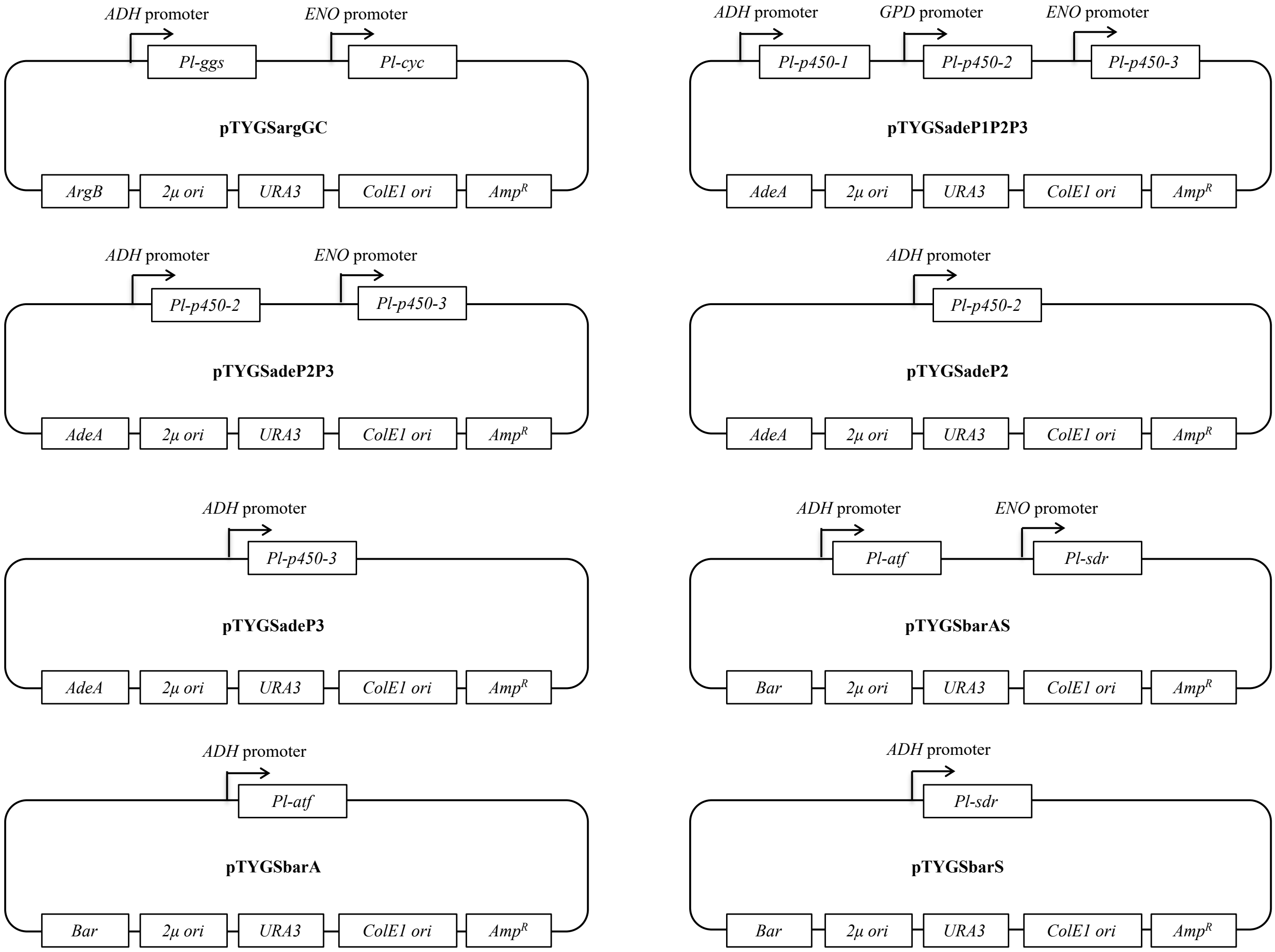

**Supplementary Figure 35.** Plasmid maps of the expression vectors used in this work.
